# Generalisation of training-induced recovery in occipital stroke: neurochemical and fMRI correlates

**DOI:** 10.64898/2026.05.29.728443

**Authors:** Hanna Willis, Lucy Starling, Isobel Rout, Brendan Sargent, Aaron Kay, Rebecca Millington-Truby, I Betina Ip, Matthew R. Cavanaugh, Sara Ajina, Krystel R. Huxlin, Marco Tamietto, Holly Bridge

## Abstract

**BACKGROUND:** Damage to the early visual cortex after an occipital stroke typically results in the loss of conscious vision in the contralateral hemifield. Nonetheless, extensive perceptual training can restore visual motion discrimination in the blind-field. Here, we assessed, in a cohort study, whether improvements transferred to an untrained Gabor detection task and whether awareness within the blind field increased. We then explored the neural underpinnings of these changes.

**METHODS:** Eighteen participants (6 female; aged 24-74 years; >6 months post-stroke) completed at least six months of visual rehabilitation in their blind field. Rehabilitation consisted of participants practicing a two-alternative, forced-choice, motion discrimination task using random dot stimuli, five days/week, at home, at one or two non-overlapping, locations in their blind-field. Each participant also completed two in-lab visits: one pre- and one post-training. A subset returned to the lab for a follow-up visit three months later to assess persistence of recovery. In addition to the trained task, an untrained, drifting-Gabor detection task was used to measure transfer of learning and changes in visual awareness at the trained locations. To investigate neural mechanisms underlying generalisation of improvements, participants completed MRI scanning at each lab visit. Magnetic resonance spectroscopy (MRS) was used to quantify GABA and glutamate concentrations in the ipsilesional motion sensitive area, hMT+, and a control voxel in the sensorimotor cortex. Functional MRI was conducted to assess BOLD signal changes in hMT+ and across the rest of the brain during passive viewing of high contrast Gabor stimuli in the blind field.

**RESULTS:** Participants showed significant improvements in motion direction discrimination (trained task) between pre- and post-training in-lab visits, which generalised to improvements in Gabor detection and awareness (untrained task). Reduced GABA and glutamate in ipsilesional hMT+ was linked to improved Gabor detection, but not awareness. Increased BOLD signals in hMT+ and dorsolateral prefrontal cortex also correlated with improved Gabor detection, while awareness changes were linked to higher-level areas associated with visual attention in the contralesional prefrontal cortex (area 46) and inferior parietal lobule.

**CONCLUSIONS:** Long-term visual rehabilitation using a global motion discrimination task generalised to enhance both detection and awareness of moving Gabors within the blind field of occipital stroke survivors. Improvements were supported by selective changes in brain regions known to be involved in motion perception and attention respectively, suggesting that a broad network supports recovery, which could be targeted to enhance outcomes.

## Introduction

Damage to occipital cortex, including primary visual cortex (V1) due to stroke leads to contralateral visual field deficits. This occurs in 10-25% of all stroke survivors^1^ and can be devastating to visual function and quality of life ^2–4^. Despite the high prevalence of visual field loss, profound consequences, and significant impact on daily life, there are currently no widely accepted rehabilitation techniques^5,6^.

Some people with damage to V1 show the ability to detect stimuli in their blind field^7–11^. Previous work using fMRI^12^, diffusion-weighted MRI^13,14^ and functional connectivity^15^ to investigate the neural basis of this residual vision (including phenomena defined as “blindsight”) has highlighted the importance of human visual motion area hMT+ and the pathway connecting it with the dorsal lateral geniculate nucleus (dLGN). Moreover, residual vision correlates with concentrations of the major inhibitory and excitatory neurotransmitters (GABA and glutamate) within hMT+^16^. Thus, evidence across multiple imaging modalities implicates area hMT+ in residual visual function.

Visual training within the blind field has been shown to restore vision in the blind field^17–23^ and, in some cases, is related to neural changes. These include increased functional connectivity in the precuneus^20^, change in population receptive field size in V1^17^ increased fMRI activity to moving stimuli in area hMT+^24^, and increased integrity of the white matter pathway between the dLGN and hMT+^14^. While these studies differed in the type of training, testing, and location of neural change, all suggest potential for increasing neural activity in the visual system of people with visual field loss.

Key to the utility of rehabilitation programmes is whether improvements generalise to other visual stimuli and tasks. In the healthy visual system, the generalisability of learning is debated, but is less likely to occur for difficult, precise tasks that require early visual processing^25^ to be altered^26^. Several studies showing visual recovery in occipital stroke patients found that improvements generalised to untrained tasks, with training on global direction discrimination and integration improving contrast sensitivity, static orientation discrimination^27^ and fine direction discrimination^28^. Additionally, training can reduce the size of clinically-assessed visual field deficits measured by light detection perimetry^19,23,29^. However, while visual restitution therapies aim to improve the lives of stroke survivors by allowing recovered vision to be used in daily life, we are still trying to understand whether restored vision is concious or unconscious (similar to the phenomenon of “blindsight”). Previous studies have shown that attentional cues assist with restitution training^28,30–32^ and therefore implies attention may be important for restoring vision.

A key question in the field of perceptual training is how generalisation of learning and recovery are related to neural changes in the brain. Relevant to the perceptual training task employed presently, global motion discrimination in primates is believed to critically depend on MT^33–36^. Thus, it is possible that when stroke survivors recover this ability after damage to early visual cortex, hMT+ neurons become more sensitive to motion stimuli generally, leading to improvements in other tasks involving motion. In the current study, we therefore expected that training for at least six months on a motion discrimination and integration task using high-contrast, random dot stimuli would enhance contrast sensitivity for detecting a moving Gabor patch. Second, we predicted that extensive rehabilitation targeting the often unattended blind field might increase awareness within this blind field. Third, we predicted that a reduction in the major inhibitory neurotransmitter GABA (measured using MRS) and increased activity (measured by fMRI) in hMT+ would be related to improved perception. Finally, we also predicted that perceived awareness in the blind field would increase with rehabilitation and would be supported by increased activity in attentional areas.

## Materials & Methods

### Study design

The study design used presently has already been detailed in one publication^14^. In summary, participants visited the Oxford Centre for Integrative Neuroimaging for up to three visits: pre-training (N=18), post-training (N=18) and follow up (N=12; Figure 1A). These visits were structured identically and invoved fundamental characterisation of each person’s visual performance as well as brain imaging. Between visit 1 and 2, participants were asked to complete at least six months of visual training at-home on a global direction discrimination and integration task (Figure 1B; minimum number of sessions = 100; median number of sessions=152.5; range=102-271) at two locations within the blind field per day (∼40minutes; 300 trials per location – see Supplementary Materials and Figure S1 for further details including how direction range was calculated and thresholds were measured). Participants returned for the post-training visit at least six months after their initial baseline visit (median±IQR=8±3 months; range=6-14 months). A subset of participants then returned after a further three months without training (median±IQR=95±15.3 days; range=84-118 days) to determine whether improvements persisted over this period. The study and planned analyses were registered on clinicaltrials.gov before the start of data collection (NCT04878861).

**Figure 1.**
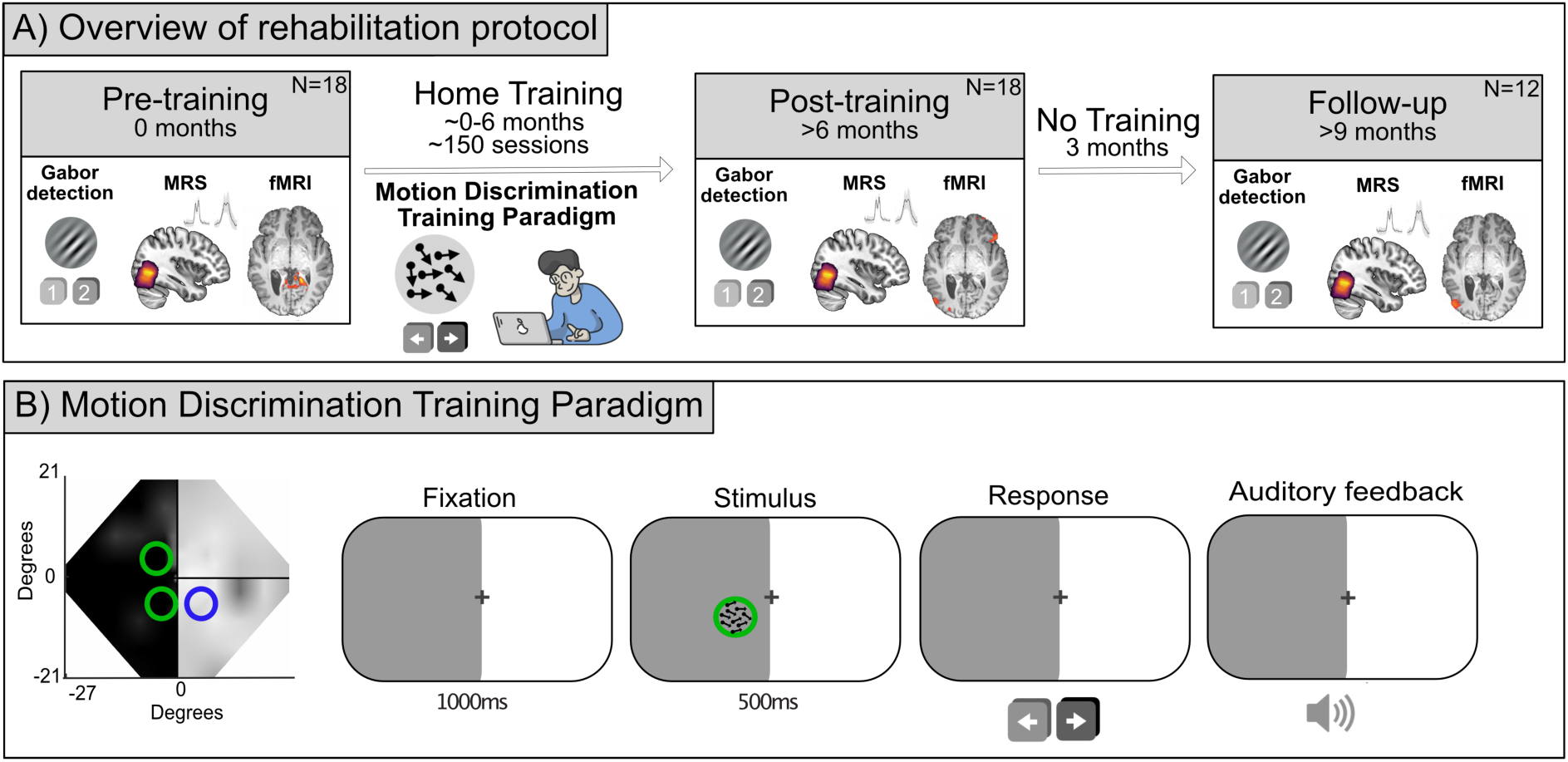
Overview of study design. A) Study design involving up to three research visits; baseline (pre-training), after at least six months (post-training) and after a further three months (follow up). Psychophysics (trained motion discrimination and non-trained moving Gabor detection), MRS and fMRI data were collected at all visits. Participants completed at least six months of visual training at two locations in their blind field between the pre- and post-training timepoints (visits 1 and 2). B) Global motion discrimination training protocol: participants began each trial by fixating centrally. A high-contrast, moving random dot stimulus was then presented at a specific training location within the blind field for 500ms. After the stimulus disappeared, participants were asked to indicate if they perceived the global direction of dot movement to be leftward or rightward. Presentation of stimuli was accompanied by an auditory tone, and auditory feedback signalled whether the response was correct or not after every trial. This was repeated 300 times for each blind field location during each training session.

### Previous publications

Pre-training behavioural^23^, MRS^16^ and structural data^37^ have already been published. Training related improvements in the direction discrimination task and changes in diffusion-weighted^14^ and structural^23^ MRI data have also been published.

### Participants

Twenty-four stroke survivors with visual field deficits were recruited. Participants were healthy, MRI-safe, English-speaking adults with damage to V1 sustained in adulthood (18+ years) that resulted in a homonymous visual field defect. Damage was sustained a minimum of 6 months prior to participation in the study. Ten age-matched controls were also included to provide concentrations of neurochemicals measured in the healthy visual system (median age=41 years; range=28-69; 4 female). Participants had no history of diagnosed cognitive or psychiatric disorders, including executive or attentional deficits. They had no history of eye disease or impairment other than visual field deficit, including all forms of visuospatial neglect.

Participants were excluded from all analyses if they did not complete the MRI scan (n=2) or they did not complete training (n=4). This left 18 participants who had MRI scans before and after six months of training (median age = 49 years; range 24-74; 6 female; full participant details in Supplementary Table S1). Of these, some were also removed from specific analyses – e.g. behavioural analyses if they did not complete the task accurately (n=1), from MRS if their data could not be fit or exceeded 3SD from the mean (n=2), and from the fMRI analyses if they did not exhibit fMRI activity to stimuli shown in the sighted field (n=3; see Supplementary Table S2 for full details). Complete multimodal datasets were therefore obtained from 13 participants; however, all participants were included in analyses for which adequate data were available.

Ethical approval was given by the University of Oxford Central Research Ethics Committee (R60132/RE001). Participants were given the study information sheet prior to enrolment and were explicitly told that the training was to inform research and could not be guaranteed to improve vision. All participants provided written informed consent, and experiments were conducted in accordance with the ethical guidelines of the Declaration of Helsinki.

### Assessing generalisation of vision restoration to a moving-Gabor detection task

Before and after global motion discrimination and integration training in their blind field, participants were tested using a drifting-Gabor detection task^12,16^. Participants were seated with a chin-forehead rest at a viewing distance of 42cm and tested at both blind field training locations and a single sighted field location. A two-interval forced-choice protocol in which participants indicated whether the Gabor stimulus appeared in the first or second interval was employed (Figure 2A). The start of the interval was indicated by a 500ms auditory tone, with a ‘low’ 300Hz tone marking interval 1 and a ‘high’ 1500Hz tone for interval 2. The stimulus appeared for 500ms with a jittered onset range of 500-1500ms while the participant fixated on a central black cross. Eye fixation was monitored throughout using an Eyelink 1000 eye tracker (SR Research Limited, Ontario, Canada). If participants moved their eyes during the task, they were reminded by the researcher to ensure they were fixating centrally. Participants were then asked in which interval the stimulus appeared (1 or 2), and whether they were aware something had appeared (yes or no) and responded with a keypress on a keyboard. Drifting Gabor stimuli (diameter=5°; sigma=0.8°; stimulus duration=500ms; spatial frequency=1c/°, temporal frequency=10Hz) were generated in MATLAB (Mathworks) and Psychtoolbox (Brainard et al., 1997) and presented on a mid-grey uniform background of luminance 16.5 cd/m^-2^ pseudo-randomly at different luminance contrasts (1, 5, 10, 50, 100%). Each contrast was repeated 10 times.

**Figure 2.**
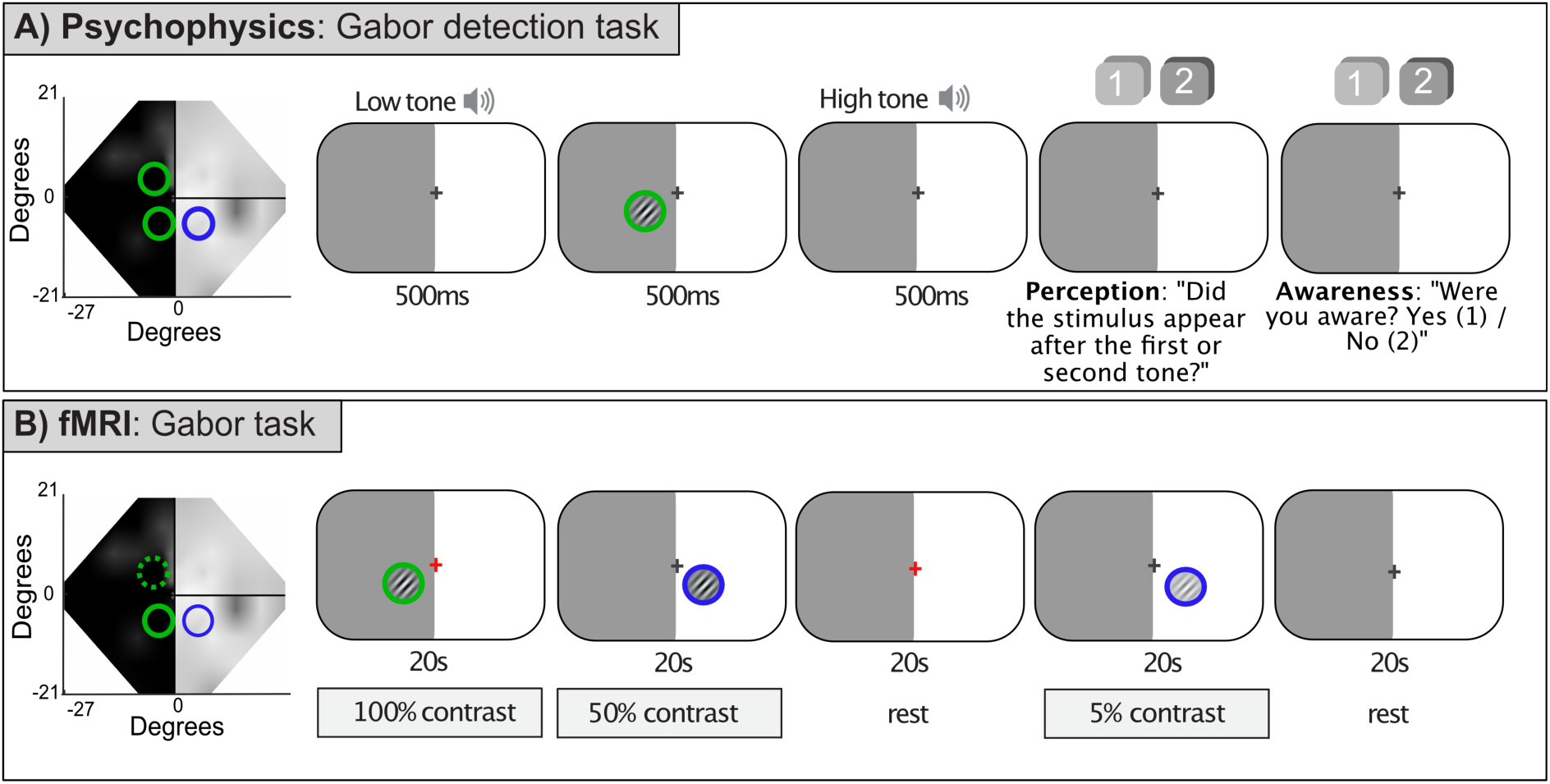
Schematic of psychophysical and fMRI experiments. The visual field loss in the left hemifield is shown on the Humphrey visual field with two example training locations (green) and sighted field location (blue) highlighted. A) The psychophysical task where a moving Gabor stimulus with varying contrast (1%, 5%, 10%, 50% and 100%) was shown in either the 1^st^ interval (indicated by a low tone) or the 2^nd^ (high tone). Participants report which interval the stimulus appeared and whether they were aware of the stimulus presentation (yes or no). This was repeated for both blind field locations and a matched sighted field. B) The fMRI experiment where moving Gabor stimuli of different contrasts (1%, 5%, 10%, 50% and 100%) were shown to one blind location and sighted field location in a block design. Participants were asked to fixate centrally and press a button whenever the central cross turned red. The blind field location where fMRI data were not collected is indicated by a dashed green line on the Humphrey field.

The percentage of trials where the participant reported that they were “aware” was taken as a measure of the participants’ awareness of the stimulus. This measure reflects self-reported visual experience and was independent of response accuracy (although see Supplementary Figure S2 for analysis of awareness for just the correct trials). Since residual vision (and “blindsight”) is seen most frequently for high contrast stimuli^12^, percentage correct and awareness for 50 and 100% contrast Gabors (Figure 3A) were also averaged separately. The visual field location at which the individual performed best with the training task was used for all further analyses (later referred to as the “most improved location”).

**Figure 3.**
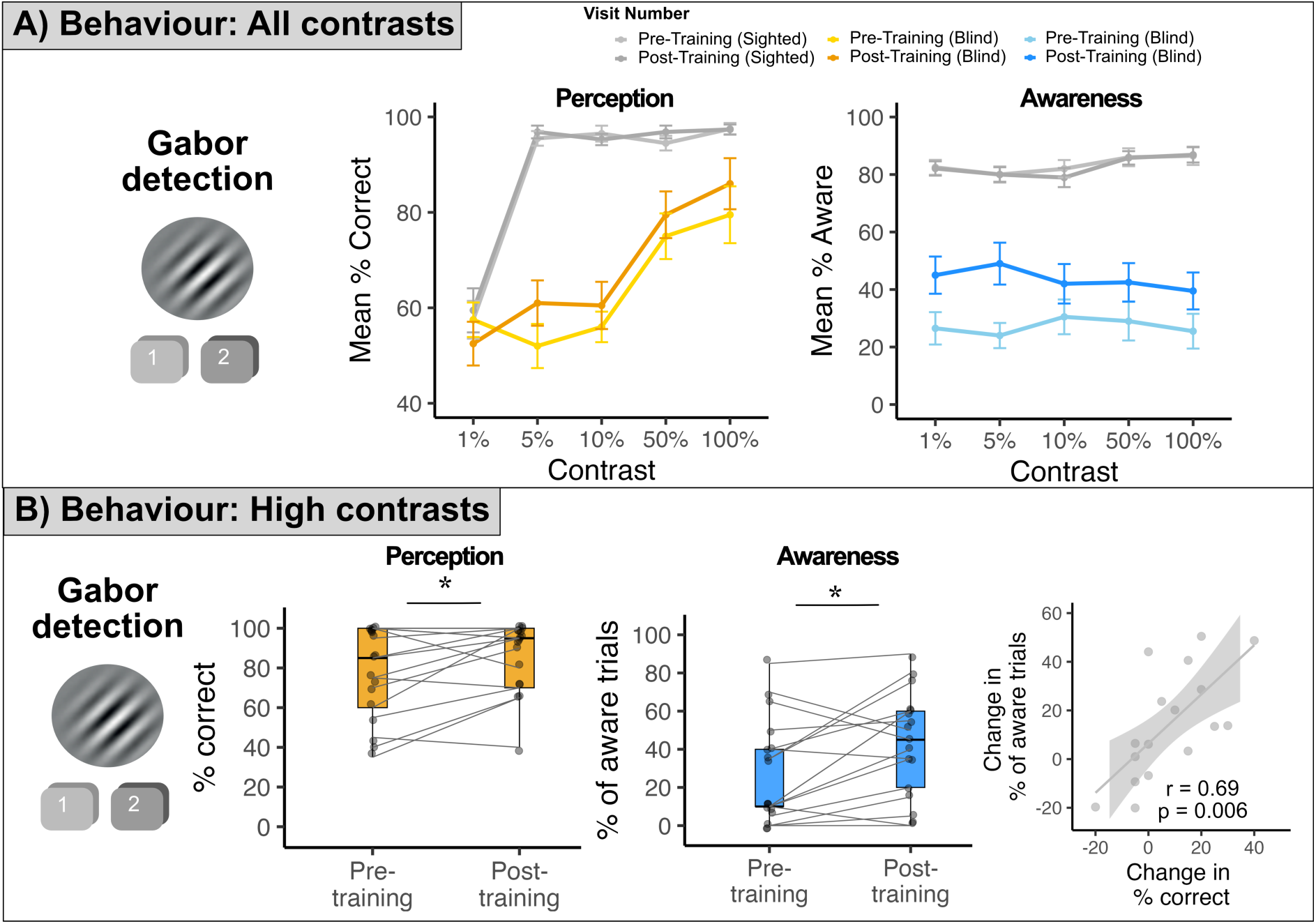
Performance changes on the moving Gabor detection task between the pre- and post-training visits. A) Shows mean percent correct (left) and percent of aware trials (right) for the blind field pre- (yellow; light blue) and post-training (orange; dark blue), and for the sighted field pre- (light grey) and post-training (dark grey) across all contrasts (1%, 5%, 10%, 50% and 100%). Since participants are only able to do this task reliably above chance for the higher contrasts, 50% and 100% contrast were combined for all further analyses. B) Shows the increase in high contrast percent correct (yellow) and percentage of aware trials (blue) between the pre- and post-training visits and the strong correlation between changes in high contrast percentage correct and changes in high contrast awareness (grey).

### fMRI acquisition

Participants were scanned using a 3T Siemens Prisma MRI scanner with a 64-channel head coil. A whole-brain T1-weighted structural image (TE = 3.97ms, TR=1900ms, FoV=192mm, flip angle = 8°) was acquired for each participant at each timepoint for registration purposes.

Stimuli for the functional MRI (fMRI) task were presented on a screen at the back of the MRI scanner bore (viewing distance=127.5cm) viewing through a front-silvered mirror. An EyeLink 1000 eye tracker (SR Research) was used to confirm central fixation. A multiband gradient echo sequence was used to acquire the fMRI data (2mm isotropic resolution, TR=1050 ms, TE=37.0 ms, flip angle=60°, multi-band factor=6).

Participants passively viewed a drifting achromatic Gabor patch (same stimulus parameters as psychophysical task described previously^12^, presented at one of the two training locations in the blind field and at the equivalent location in the sighted field, in a block design (see supplementary materials for details).

### MRS acquisition

MRS data were acquired from two separate voxels (25x25x20mm^3^), one placed over hMT+ and one placed over the sensorimotor cortex (SMC) in the ipsilesional hemisphere (see Supplementary Materials for positioning and repositioning details) at each timepoint. A locally developed MEGA-PRESS (MEscher-GArwood Point RESolved Spectroscopy^38^ derived from the CMRR spectroscopy package MEGA-PRESS sequence (https://www.cmrr.umn.edu/spectro/) was used to measure GABA+ and Glx (see Supplementary Materials for specific details).

### fMRI analysis

Preprocessing and all whole-brain statistical analyses were carried out using FSL tools (version 6.00 www.fsl.fmrib.ox.ac.uk; see Supplementary Materials for pre-prcocessing details). To perform group analyses, all brains where the lesion was in the left hemisphere were flipped so that the right hemisphere was always the ‘lesion’ side. Using FMRIB’s easy analysis tool (FEAT) each different contrast level and hemifield was included as an explanatory variable in a general linear model and compared to rest (fixation without a stimulus). Additional contrasts included ‘high contrast’ which included trials with both 50% and 100% contrast.

Group level analyses were performed using both mixed-effects and fixed-effects analyses. While mixed-effects approaches account for inter-subject variance, allowing for generalisation to a wider population, the relatively small sample and variability in both the cortical damage and BOLD signal mean that this is a conservative measure. A fixed-effects analysis is valid for the specific population studied, and activation was only considered in the occipital lobe where there were pre-existing hypotheses about changes in activity following training. The main paired t-test contrasted the activation post-training compared to pre-training across participants. A comparable paired t-test was also performed between the post-training and follow-up sessions to determine whether any changes persisted beyond the end of training. A second group approach identified BOLD signal changes between post- and pre-training that correlated with the change in performance on the psychophysical Gabor detection task, used as a behavioural covariate. MRI images are shown on MNI standard brains in MRIcroGL^39^ (version 14.5).

### MRS analysis

As with previous publications^16^, the MRS data for both voxels (one over hMT+ and one over sensorimotor cortex) and each timepoint (pre-training, post-training and follow up) were analysed using a local version of Gannet 3.1, an open-source toolbox for analysing edited MRS data (http://www.gabamrs.com/; see Supplementary Materials for analysis details). MRS voxels at each timepoint were reconstructed using FSL-MRS^40^ (Version 2.4.0). See Supplementary Materials for analyses showing similar voxel positioning across timepoints (Figure S1) and no differences in fit error or full-width half-maximum (FWHM) between timepoints in either voxel of interest or between patients and controls (Tables S3 and S4).

### Statistical analysis

All statistical analyses were carried out in R studio (R version 4.1.2) using the *rstatix* package. Data are presented as median *±* interquartile range (IQR) unless otherwise indicated. Where data met the assumptions, parametric tests were used, otherwise non-parametric tests were used. Paired tests (either t-tests or Wilcoxon tests) were used to compare timepoints for the Gabor detection task and MRS. Effect sizes are reported for parametric tests (Cohen’s d) and for non-parametric tests (r). All reported p values were Holm-Bonferroni corrected.

## Results

### Global motion training generalises to improve detection and awareness of moving Gabors

As previously reported, global motion and integration training significantly improved performance on the trained task at the trained, blind-field locations^14,23^. Figure 3A presents the Gabor detection data for the 17 participants who completed both pre- and post-training sessions, shown at each individual’s “most improved” blind field (yellow; blue) location and a matched sighted field location (grey) across all contrast levels.

Since residual vision is most evident for high contrast stimuli (evident in Figure 3A where 1%, 5% and 10% are all at chance), 50% and 100% contrast conditions were and used for all subsequent analyses (Figure 3B). Between the pre- (N=17; median±IQR=80±35%) and post-training visits (N=17; median±IQR=95±25%), there was a significant improvement in detection of high-contrast, drifting Gabors (Figure 3B yellow; paired t-test: t(16)=-2.19; p=0.044; Cohens d=0.38). This was associated with a significant increase in the percentage of aware trials between pre- (Figure 3B blue; N=17; median±IQR=20±30%) and post-training visits (median±IQR=45±37.5%; paired t-test: t(16)=-2.66;p=0.034; Cohen’s d=0.54). Additionally, there was a strong correlation between percentage correct performance and awareness at both pre- (Pearson’s correlation: r=0.79; p<0.001) and post-training visits (Pearson’s correlation: r=0.79; p<0.001; see Supplementary Figure S2), as well as between *change* in percentage correct and *change* in awareness following training (Figure 3B grey; Pearson’s correlation: r=0.69; p=0.006). In sum, participants showing greater improvements in perception, also showed greater increases in awareness of the stimuli in their blind field. There was also a significant increase in the proportion of correct responses of which the participants reported being aware (see Supplementary Figure S3 and associated text).

Surprisingly, there was no correlation between the change in performance in the trained (motion discrimination) and untrained (Gabor detection) tasks: there was no significant correlation between normalised direction range threshold change and percentage Gabor detection correct change (Pearson’s correlation: r=-0.08; p=0.77) or percentage of aware Gabor trials (Pearson’s correlation: r=-0.01; p=0.96).

### No group-level change in neurochemistry or BOLD signals in hMT+ following 6 months of training

To determine whether training induced neurochemical changes in participants’ lesioned hemispheres, GABA+/tCr and Glx/tCr concentrations were quantified in voxels placed over ipsilesional hMT+ and in a control voxel in the ipsilesional SMC. As can be seen in Figure 4A, values for GABA+/tCr in the hMT+ voxel were similar at the pre-training (n = 16; 0.11 ± 0.04; median±IQR) and post-training (n = 16; 0.10 ± 0.02) visits, with no difference between concentrations at the two timepoints (paired t-test:GABA+/tCr; t(15)=1; p=0.33). Similarly, for Glx, the average concentration was the same pre- (n = 16; 0.07±0.02) and post-training (n = 16; 0.07±0.02), with no difference between the timepoints (paired Wilcoxon: W = 144; p = 0.56). These values were comparable to GABA+/tCr (0.10 ±0.02) and Glx/tCr (0.06 ±0.01) measured in hMT+ in neurologically healthy age-matched participants. There was no difference in GABA+/tCr (independent t-test: t(20.88)=1.21; p=0.24) or Glx/tCr (independent t-test: t(19.52)=1.19; p=0.25) in hMT+ between stroke survivors (at the baseline visit) and controls. As can be seen in Figure 4B, there was also no evidence of BOLD signal change in hMT+ after rehabilitation when using a whole brain analysis for either fixed or mixed effects.

**Figure 4.**
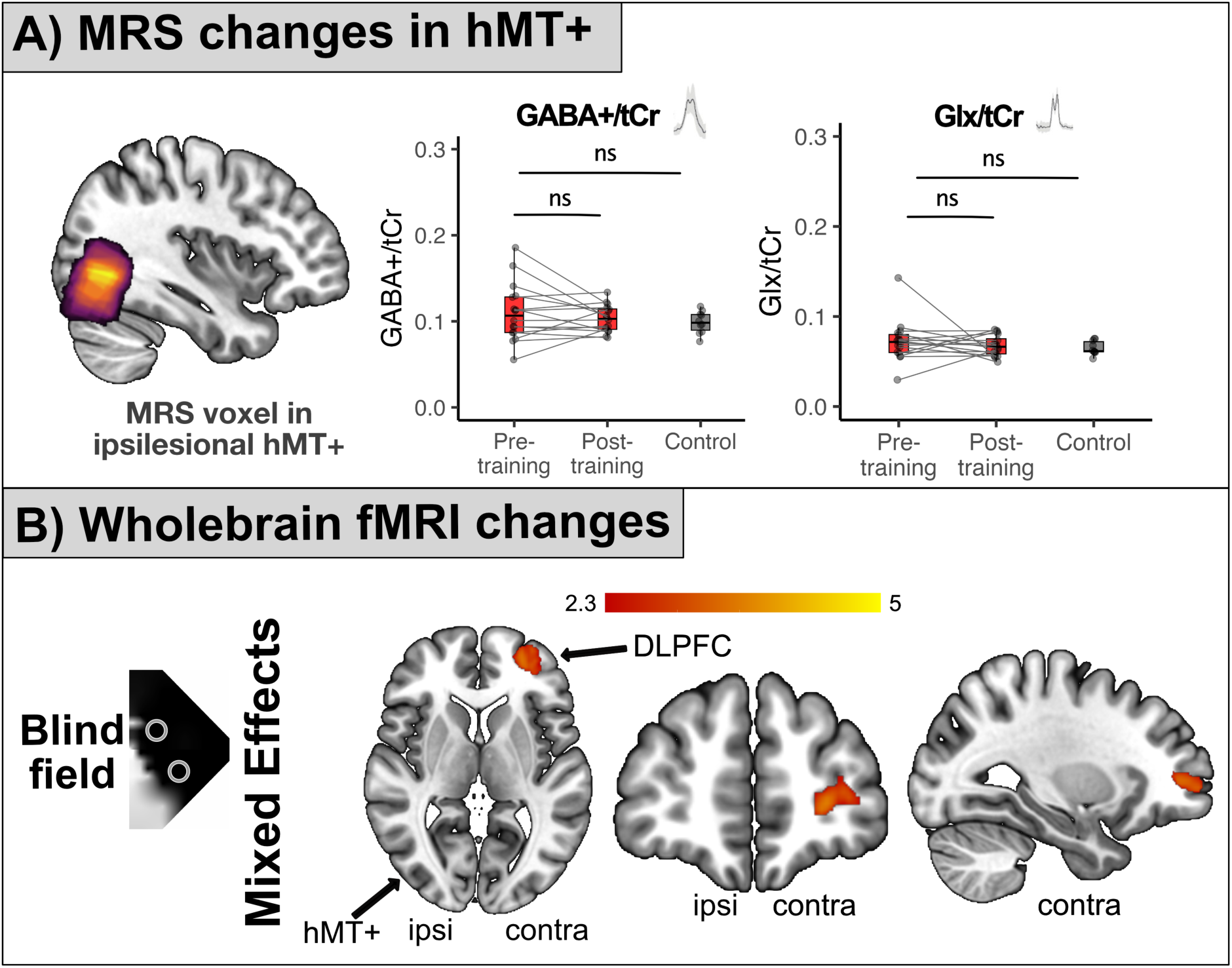
Neural changes between pre- and post-training visits. A) The overlap of hMT+ voxel across participants in the ipsilesional hemisphere (left) and the concentrations of GABA+/tCr and Glx/tCR in hMT+ (right). Values for participants with visual field deficits (red) are comparable to healthy controls (grey), and there was no significant difference between pre- and post-training values of GABA+/tCr or Glx/tCr in hMT+ (ns) or between stroke survivors at the pre-training visit and controls (ns). B) The increase in neural activation to high-contrast stimuli in the post-training compared to the pre-training scan. Using a mixed-effects analysis, the dorso-lateral prefrontal cortex showed a significant change to blind field stimulation while no changes were found in hMT+. Ipsilesional (ipsi) and contralesional (contra) hemispheres are indicated.

The control SMC voxel showed no difference in GABA+/tCr (pre-training = 0.13±0.03; post-training = 0.12±0.04; paired t-test: t(15)=1.64; p=0.0.12) or Glx (pre-training = 0.07±0.02; post-training = 0.07±0.02; paired Wilcoxon; W=124; p=0.90) concentration across the two time points. Healthy control participants had a similar concentration of GABA+/tCr (n=10; 0.11±0.04; independent t-test: t(23.17)=0.91; p=0.37) and Glx/tCr (n= 10; 0.07±0.03; independent Wilcoxon; W=86; p=0.78) concentration in SMC compared to stroke survivors (at the baseline visit).

It should be noted that although quantifications of GABA/tCr (∼0.10) are comparable to previous literature (∼0.10-0.15)^41,42^, Glx/tCr (∼0.07) values were slightly lower than past literature (∼0.08-0.12)^43,44^. However, they were comparable across all participants and voxel locations which suggests the differences were likely due to methodological or modelling differences rather than reductions in stroke survivors.

### Increased BOLD signals in dorso-lateral prefrontal cortex after 6 months of training

To determine whether there was any change in neural activity due to training, a whole-brain paired t-test was performed across the 15 participants who showed consistent fMRI activity to stimuli presented in the sighted field. Figure 4B shows the regions with a significant increase in response following the training. The results were identical across fixed and mixed effects analyses so only the conservative mixed effects analysis is presented here (see Figure S5 in the supplementary materials for the fixed effects results). The only region that showed a significant increase in both analyses was the dorso-lateral prefrontal cortex (DLPFC) in the hemisphere contralateral to the training (and ipsilateral to the lesion). Notably, this increased activation was present for stimuli shown in the blind and sighted visual hemifields (see Figure S6 in in the supplementary materials for the sighted field results).

### Persistence of behavioural and neural changes

Twelve participants returned three months after their post-training visit to determine whether the behavioural and neural changes persisted. Eleven participants were included in this analysis, as R010 was removed (see Supplementary Materials for details). Between the post-training (N=11; median±IQR=80±30%) and follow-up (N=11; median±IQR=90±15%) sessions there was no significant change in percentage correct in the Gabor detection task (Figure 5A; paired t-test: t(10)=-1.13; p=0.28), suggesting that participants maintained their improvements. Moreover, their awareness also did not differ between the post- (N=11; median±IQR=35±37.5%) and follow up (N=11; median±IQR=35±25%) visits (Figure 5A, middle panel; paired t-test: t(10)=-0.76; p=0.46). However, perceptual and awareness measures were no longer correlated at the follow-up timepoint (Pearson’s correlation; r=0.54; p=0.08), although a trend remained. We posit that the loss of significant correlation is potentially due to the reduced sample size in this analysis.

**Figure 5.**
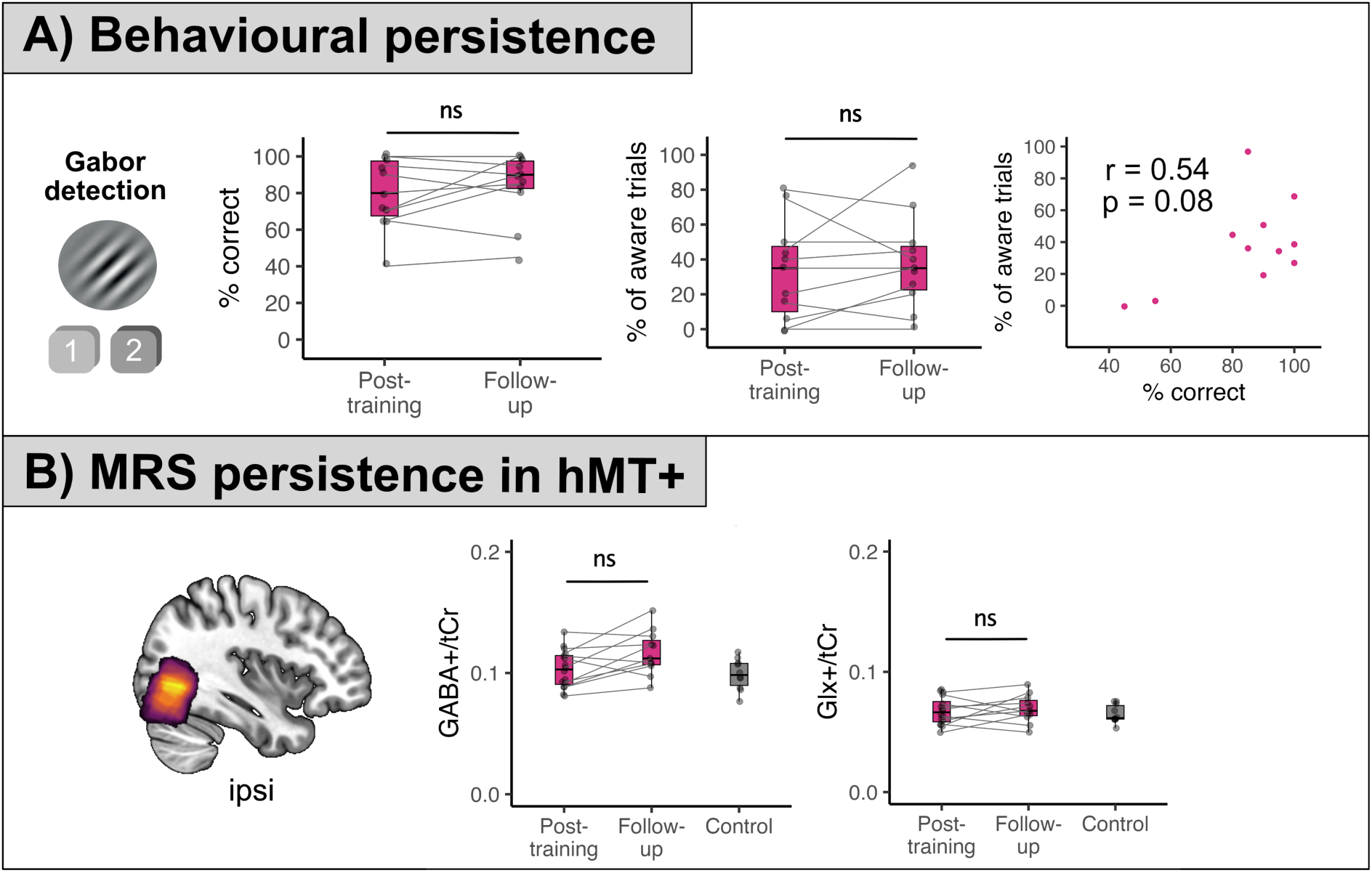
The persistence of behavioural and neural changes following 3 months without training. A) No significant change (ns) in performance on the Gabor detection task (left) or stimulus awareness (middle), and a lack of correlation between high contrast Gabor detection performance and awareness (right). B) Shows the neurochemical concentration of GABA+ (left) and Glx (right) at post-training and follow-up, which were not significantly different.

Since there was no change in neurochemical concentration in hMT+ as a result of training, it is not surprising that there was no change following cessation of training. The concentration of GABA+/tCr in hMT+ (Figure 5B) did not differ between the post-training (median±IQR=0.10±0.02) and follow-up visit (median±IQR=0.11±0.02; paired Wilcoxon: W=10; p=0.08). There was also no significant change in Glx/tCr in hMT+ (Figure 5B; post-training median: 0.07; follow up median=0.07; paired Wilcoxon: W = 24, p = 0.46). Unsurprisingly, in SMC, no changes were noted between post-training and follow-up visits for either GABA+/tCr (post-training median±IQR=0.11±0.04; follow up median±IQR=0.12±0.02; paired Wilcoxon: W=30, p=0.83) or Glx/tCr (post-training median±IQR=0.12±0.04; follow up median±IQR=0.12±0.02; paired Wilcoxon: W=56, p = 0.08).

Finally, using a mixed-effects analysis, we observed no changes in neural activity between post-training and follow-up fMRI scans. The fixed-effects analysis, which only considered changes in the occipital lobe, did not show any reduction in neural activity at follow-up, but there was an increase in occipital activation compared to the post-training session. Specifically, there was some increased activation in ipsilesional ventral visual cortex and an increased in contralesional hMT+ activation to blind field stimulation (see Figure S7 in the Supplementary materials). However, due to the reduced sample size of this analysis, these results should also be considered with caution.

### Neural mechanisms correlating with change in perception

We next considered if the lack of group difference in neurochemistry or neural activity in the ipsilesional visual cortex following training might be related to variability in the magnitude of visual improvement across individuals. Indeed, we found a significant correlation between improvement in Gabor detection ability (measured by percent point increase) and change in GABA+/tCr in hMT+ (Pearson’s correlation; r=-0.60; p=0.04); the greatest performance improvements were associated with the greatest decreases in GABA+/tCr (Figure 6A left). Similarly, improvement in percentage correct was inversely correlated with a reduction in Glx/tCr in hMT+ (Pearson’s correlation; r=-0.58; p=0.04; Figure 6A right). No such correlations were observed for the control voxel in SMC (GABA+/tCr: Pearson’s correlation; r=-0.16; p=0.55; Glx/tCr: Pearson’s correlation; r=0.06; p=0.81).

**Figure 6.**
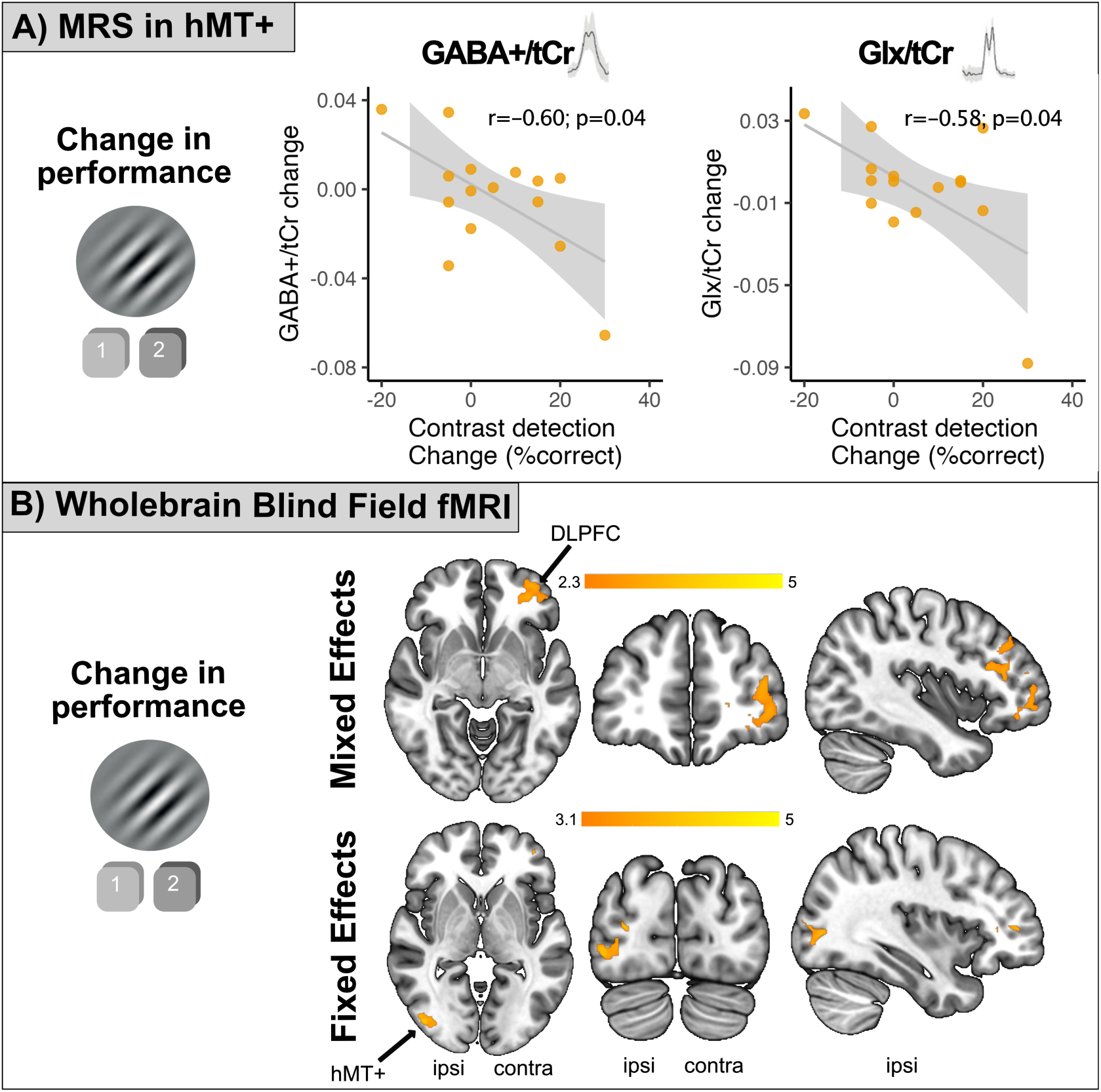
Neural correlates of change in Gabor performance. A) Shows the significant negative correlation between the change in Gabor detection performance and GABA+/tCr concentration in hMT+. A similar correlation was found with Glx/tCr indicating that improved performance was related to decrease in both GABA+/tCr and Glx/tCr after training. B) Shows the regions of the brain where the increase in performance was correlated with increased BOLD signal change to blind field stimulation. The mixed-effects analysis shows regions of the DLPFC, including areas 46 and 9 in the hemisphere contralateral to the damage. The fixed-effects analysis indicated that for this population, hMT+ in the lesioned hemisphere also showed an increase in activity that correlated with the improvement in visual performance. “Ipsi” and “contra” indicate the ipsilesional and contralesional hemispheres respectively.

To determine whether changes in fMRI activity following training were also related to the extent of improvement in visual function, the same percent point change in the Gabor detection task was used as a correlate. Figure 6B shows brain regions whose activity increased when visual performance improved following training. The mixed effects analysis revealed that in the DLPFC, including both area 46 and 9, the increase in activation correlated with improved performance. The less conservative fixed-effects analysis indicated that change in activity in area hMT+ (black arrow) was also significantly correlated with improvement in performance.

A similar analysis was performed to determine the relationship between change in normalised direction range thresholds on the motion discrimination task and neural measures. The DLPFC also showed an increase in activity that correlated with this measure of visual performance. See Supplementary Figure S4 for details.

### Neural substrates of increased blindfield

Lastly, we asked whether neurochemistry in hMT+ or BOLD signal change across the brain was related to improvement in subjective awareness within the blind-field, across participants. We found no relationship between improvements in awareness on the Gabor detection task and changes in neurochemistry, in either GABA+/tCr (Pearson’s correlation; r=-0.07; p=0.80) or Glx/tCr (Pearson’s correlation; r=-0.06; p=0.83) in hMT+ (see Figure 7A). However, due to the high correlation between percentage correct and awareness, potential effects may have been obscured. To disentangle their individual contributions, a multiple linear regression was conducted to assess the effects of both awareness and percentage correct on GABA levels, while controlling for each other (model: GABA+ ∼ awareness + percentage correct). The regression model significantly predicted GABA/tCr levels (F(2, 12) = 4.33, p = 0.038). Notably, percentage correct was a significant predictor (p = 0.016), whereas awareness change was not (p = 0.284). Additionally, no relationship was found between awareness and GABA+/tCr (Pearson’s correlation; r=0.29; p=0.26) or Glx (Pearson’s correlation; r=0.28; p=0.28) in control voxel SMC.

**Figure 7.**
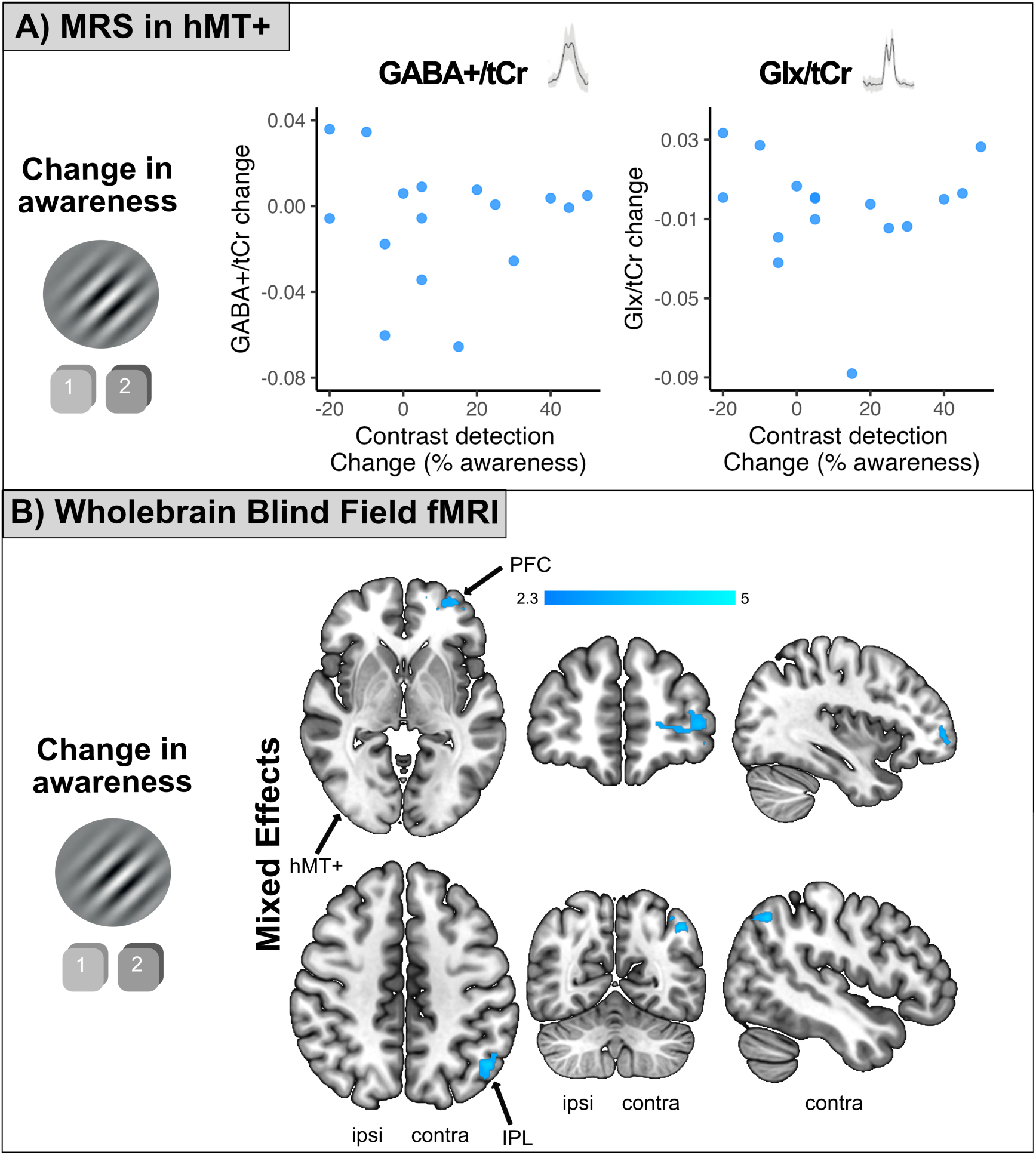
Neural correlates of change in awareness. A) Shows no correlation between the change in awareness and change in GABA+/tCr or Glx/tCr concentration in hMT+ after training. Each datapoint represents the data from an individual participant. B) Shows the regions of the brain where the increase in awareness was correlated with increased BOLD signal change across the group to blind stimulation after training. The mixed-effects analysis shows regions of the prefrontal cortex (area 46) and the inferior parietal lobule in the hemisphere contralateral to the damage. “Ipsi” and “contra” indicate the ipsilesional and contralesional hemispheres respectively

Using a mixed-effects analysis, the increase in awareness measured in the behavioural task was significantly correlated with an increase in fMRI activity in the prefrontal cortex (area 46) of the non-damaged brain hemisphere when stimuli were presented to the blind field (see Figure 7B). When a fixed effects analysis (valid only for the current population) was used, activity was seen in bilateral PFC and IPL (see Figure S8 in Supplementary Materials). There was no evidence of a relationship between activity in hMT+ and awareness across either analysis. Change in activity in the inferior parietal lobule was significantly correlated with the increase in awareness only when stimuli were presented to the blind field.

## Discussion

In this visual rehabilitation study, at least six months of visual training within the blind field led to improved perception for a moving Gabor detection task, accompanied by an increase in reported awareness, which persisted for 3 months beyond the end of training. This visual recovery correlated with increased visually-evoked BOLD activity in dorso-lateral prefrontal cortex and ipsilesional hMT+. Moreover, improved task performance also correlated with a reduction in GABA+ and Glx in ipsilesional hMT+, with participants who displayed greater improvements in the Gabor detection task having a greater decrease in GABA+ and Glx at the post-training visit. Finally, improvements in subjective awareness were unrelated to neurochemistry in hMT+ but were related to increased activity in several attention-related visual areas.

### Neural changes after visual rehabilitation

In the present study, we observed increased activity in the dorsolateral prefrontal cortex (DLPFC) following rehabilitation. Notably, improvements on the Gabor detection task were correlated with enhanced activity in both the contralesional DLPFC and ipsilesional hMT+, aligning with prior findings that rehabilitation can increase activity in hMT+^18^. These results additionally suggest that attentional networks may play a role in visual recovery.

Supporting this idea, training on a motion discrimination task in healthy participants was reported to increase bilateral DLPFC activity, which correlated with behavioural improvements^45^. The dorsolateral prefrontal cortex (area 46) is known to be play a key role in attention by delivering top-down signals that enhance activity in sensory cortices, with lesions in these areas leading to increased distractibility (see Szczepanski & Knight^46^ for a review). After occipital stroke, patients are often encouraged to rely on compensatory strategies that ignore the blind field^6^. However, intensive visual training may instead draw attention toward the blind field, facilitating the use of residual visual functions or “blindsight”.

Additional evidence underscores the importance of attention in visual rehabilitation. Pre-attentive cues have been shown to enhance learning transfer to untrained tasks^31^ and untrained visual field locations^32^. Moreover, hMT+ activity is known to be modulated by attention^47^, suggesting that directing attention toward motion stimuli in the blind field could boost activation in motion-sensitive extrastriate areas. Together, these findings highlight the potential role of attentional mechanisms in supporting visual recovery following stroke.

### Role of neurochemistry in plasticity

Another finding of the present study was that change of GABA+ and Glx in hMT+ (but not SMC) was related to improvement in perception in stroke survivors with visual field loss. This fits with our previous work which showed that lower GABA+ and Glx were related to better residual vision in this patient group^16^. Similarly, in chronic motor stroke survivors, decreases in GABA+ were related to improvements in rehabilitation^48^. This aligns with other studies showing that GABA reduction relates to plasticity in the healthy visual system^49^. Therefore, reduced GABAergic activity in hMT+ may help support neuroplastic changes necessary for visual rehabilitation in occipital stroke. Previous research in healthy individuals has shown that low GABA in the occipital lobe aids target detection in clutter, while higher GABA enhances fine feature discrimination^50^. Since the V1-damaged visual system appears to suffer from increased levels of internal processing noise^28^, reduced GABA in hMT+ after rehabilitation may correspond to improved target detection resulting from boosting the signal-to-noise ratio.

### Improvements in awareness supported by higher-level attention networks

While improved visual performance is clearly an important outcome of long-term training, increasing awareness to visual stimuli in the blind field is also extremely advantageous. In the current study, awareness was based on subjective report rather than accuracy-conditioned performance. An increase in awareness reflects a change in perceived visibility that may include both correct and incorrect trials. However, this study also found a highly significant correlation between visual performance and awareness. Typically, “blindsight” or residual vision after damage to V1 is the ability to discriminate visual stimuli without awareness and participants respond by guessing^10^. However, the correlation between perception and awareness observed pre-training here suggests that the classical definition of blindsight may not reflect the experience of the majority of participants in this study, although awareness always remains low (less than 50% of trials even after rehabilitation, see Figure 3) and is similar to previous work measuring subjective confidence^12^. The 6-month training programme used presently also increased the number of trials where participants reported being “aware” of the contrast-defined stimuli, consistent with a previous study that found visual training to increase stimulus awareness^51^.

We additionally found that improvements in awareness were related to increased BOLD signal in regions of the prefrontal cortex (area 46) and the inferior parietal lobule in the hemisphere contralateral to the damage. The prefrontal and parietal cortices are known to be involved in attention (see ^46^ for a review), with studies in humans and non-human primates showing them to be involved in top-down attention switching for visual tasks^52^. It is therefore possible that as participants increasingly deployed attention towards their blind field with visual training, this caused an increase in saliency (or awareness) of stimuli.

However, it should be noted that the measure of awareness used here was very basic, requiring participants to simply respond “yes” or “no” as to whether they were aware of stimuli. This method depends on all participants interpreting the question in the same way. Participants were encouraged to respond “yes” if they saw something, whether or not it looked like the expected stimulus. Therefore, in future studies, it would be worth using alternative awareness probes and metrics to provide additional resolution for people’s subjective experience of the presented tasks and stimuli.

### Improvements in a Gabor detection task persist without continued training

In the healthy visual system, perceptual learning for motion discrimination tasks can last for days^53^ and even months^54–56^ following the end of training. We have recently shown that the same cohort of participants as is studied here have persistence of improvement for the trained task three months after completion of training^23^. We now show that improvement in a Gabor detection task, and awareness for this stimulus, also persists beyond the training period, with the majority of participants maintaining considerable improvement from baseline. It is reassuring that persistence is maintained across both trained and transfer visual tasks, and encouraging for future rehabilitation studies.

### Limitations and future directions

A significant challenge in stroke research is variability among stroke survivors, small sample sizes, and dropout throughout a study. Damage can differ significantly even within the same brain region (e.g. occipital cortex) and is influenced by factors like age and stroke type. Therefore, due to the relatively low sample size (N=24), it was not possible to control for stroke type or location, age or time since the stroke. Despite this, recruitment of 24 people with occipital stroke is larger than many previous studies^18,19,21^, although dropout meant that MRI data were only available for 18 (75%) of these participants. While this can provide a good indication of the effects of training, variability in stroke location and size is likely to have weakened measured change in neural activity at the cohort level. Thus, a fixed-effects analysis had to be applied to the fMRI data to see the increase in activation in hMT+ that correlated with visual improvement.

Additionally, although MRS is a useful technique for measuring neurochemicals in vivo, there are limitations that should be considered when interpreting the relationship between neurochemicals and behaviour. First, GABA-editing sequences from 3T MRI scans result in a combined measure of Glx (glutamate and glutamine), and the GABA signal is affected by macromolecules and homocarnosine. Moreover, MRS provides an average measurement of neurochemical concentrations over a large cortical area (2×2×2cm) and time frame (∼8minutes); this does not allow for direct measurement of synaptic activity or specific neurochemical distribution within the voxel. Finally, MRS analyses often use a reference neurochemical, such as creatine or water, for quantification. In this study, GABA+ and Glx signals were referenced to creatine, as in an earlier study in this patient group^16^, and because we previously found that GABA+ and Glx signals referenced to water were significantly correlated with the degree of overlap between the stroke-induced fluid-filled zone and the MRS voxel^16^. Based on this, creatine was determined to be the more appropriate reference metabolite for this patient group.

Lastly, the study was designed to determine the extent of visual improvement resulting from a six-month training programme that could be feasible to implement within a clinical setting. Nonetheless, as might be expected with this type of longitudinal study, there remained considerable variability in the amount of training that each participant performed (between 6-14 months) and the time period between pre- and post-training sessions. Moreover, the amount of training required to see a modest improvement in visual performance (more than six months) remains a challenge to making this type of visual training widely available. Furthermore, despite the improvement in task performance post-training, awareness remained relatively low (∼30% trials), suggesting that there is considerable scope to increase this ability, allowing individuals to more optimally use their recovered vision in daily life. Clearly, more work is needed to boost multiple aspects of recovery, including finding approaches, such as brain stimulation or more engaging training, that can shorten training duration.

## Conclusion

The present study reports that training on a motion discrimination task transfers to a moving-Gabor-detection task and increases awareness for this untrained stimulus in the blind field of chronic occipital stroke patients, which in most people, persisted beyond the training period. The improvement in perception was accompanied by an increase in the neural activity in ipsilesional hMT+, which may be driven by changes in its neurochemistry facilitating greater plasticity. In contrast, improved awareness corresponded to increased activity in higher-level attention networks. Overall, our fidings suggest that both sensory and cognitive reorganisation are engaged in rehabilitation for stroke-induced vision loss.

## Acknowledgements

We would like to thank the participants who generously gave up their time for this research. We also thank radiographers Jon Campbell, David Parker, Michael Sanders, Nicola Aikin, and Juliet Semple, optometrist Patsy Terry, and optometric technician Charlene Hennesey, for their assistance with training and data collection.

## Data availability

Behavioural data and analysed MRS data will be shared openly on OSF. All scripts used for analysis will be shared on Github. MRI data will be made available on request.

