## Supplementary Figure 1 for "Generalisation of training-induced recovery in occipital stroke: neurochemical and fMRI correlates"

### **Supplementary materials**

#### Participants

Supplementary Table 1. All participant demographics.

| Subj | Stroke Cause | Deficit Side | Age | Time since lesion (Months) | Rehab length (Months) | Length to follow up (Days) | No of training sessions |
| --- | --- | --- | --- | --- | --- | --- | --- |
| R002 | unknown | right | 21-30 | 39 | 8 | 98 | 173 |
| R003 | ischaemic | both | 31-40 | 58 | 7 | 115 | 173 |
| R004 | haemorrhagic | left | 51-50 | 31 | 8 | 89 | 120 |
| R005 | haemorrhagic | left | 61-70 | 21 | 7 | 90 | 111 |
| R006 | ischaemic | left | 21-30 | 13 | 7 | 87 | 131 |
| R007 | haemorrhagic | left | 41-50 | 7 | 7 | 100 | 164 |
| R010 | ischaemic | right | 61-70 | 47 | 7 | 93 | 156 |
| R011 | unknown | left | 61-70 | 7 | 13 | 115 | 160 |
| R013 | haemorrhagic | right | 71-80 | 26 | 7 | 97 | 166 |
| R014 | ischaemic | right | 31-40 | 30 | 14 | 118 | 154 |
| R015 | ischaemic | right | 31-40 | 29 | 10 | 84 | 196 |
| R017 | ischaemic | right | 61-70 | 6 | 11 | - | 271 |
| R018 | ischaemic | right | 31-40 | 26 | 9 | - | 151 |
| R019 | ischaemic | left | 41-50 | 31 | 7 | 86 | 148 |
| R020 | unknown | left | 41-50 | 297 | 11 | - | 102 |
| R022 | ischaemic | right | 41-50 | 14 | 10 | - | 144 |
| R023 | unknown | right | 61-70 | 49 | 9 | - | 108 |
| R024 | ischaemic | left | 51-60 | 13 | 7 | - | 118 |
|  |  | **Median** | **47** | **27.5** | **8** | **95** | **152.5** |

#### Home training

Each participant was provided with a chin-forehead rest and software customised to their own computer and monitor specification (dimensions, resolution, refresh rate). Full instructions to set up and run the training were provided to participants at their pre-training visit. Participants performed one session at each training location each day. Data were analysed separately for the two locations. Participants were required to determine whether the global motion of black random dots on a grey background was to the left or right. Difficulty was modulated with a 3:1 staircase; after 3 correct responses the direction range increased in steps of 40° from 0° (all dots moving in the same direction) to 360° (random movement) and after one incorrect response it decreased by 40°. Auditory feedback was provided for each trial. A Weibull function was fitted to the data with a criterion threshold of 72% correct. The session threshold was normalised to the maximum range of dot directions (360˚) to generate a normalised direction range (NDR) using the following equation:

NDR threshold (%) = (360˚–Weibull-fitted direction range threshold)/360˚ × 100

For all training locations, the full diameter of the stimulus was inside the blind field border, defined at baseline with a clinical visual field test (areas with <10dB), the Humphrey visual fields. During at-home training, as performance at each location reached 72% correct and NDR thresholds stabilised over 5-10 consecutive sessions, this training location was moved to a new location 1 degree further into the blind field, laterally along the x-axis (Cartesian coordinate space), and daily training started anew.

Participants were asked to fixate centrally throughout home training. They were reminded that fixation was essential to ensure the visual target was presented at the intended training locations in their blind field. Prior experience with this approach has shown that participants are highly motivated and compliant^1,2^. An Eyelink 1000 Plus eye tracker (SR Research Limited, Ontario, Canada) was used at all assessment sessions to verify fixation and ensure fixation-contingent stimulus presentation in the lab. Only after at-home training results and threshold improvements were verified in-lab with controlled fixation, were specific participants classified as having experienced visual improvement at their training locations.

#### MRI acquisition

##### fMRI task

During the fMRI task, in each 20s block, a single contrast (1%, 5%, 10%, 50%, 100%) or rest block was presented, with the stimulus presented 8 times (2-s duration, interstimulus interval of 500ms) in each contrast block. Each participant completed three runs with the total scan time being ~11 minutes. Participants were required to maintain fixation on a static cross and press a button when the colour of the cross changed from black to red. Colour changes were random and lasted for 300ms. In three cases, the visual field loss was in the far periphery. The fixation cross was then positioned eccentrically to enable the Gabor stimulus to be presented in the blind region within the field of view available inside the scanner.

##### MRS acquisition

For the MRS acquisition the following parameters were used: TR=1500ms, TE=68ms, 160 edit-on and 160 edit-off per condition, VAPOR and dual-band editing pulse water suppression, 22.3ms editing pulse using a 53Hz bandwidth, which was centred at 1.9 ppm (“on” condition) and at 7.5 ppm (“off” condition) in alternating transients; 16-step phase cycling. For each voxel of interest, a single transient was collected with water suppression disabled (TR=2500ms). During acquisition GRE shimming was used to ensure that the water- unsuppressed MRS-measured FWHM were <12Hz and vendor-reported full-width-at-half-maximum (FWHM) were below 20Hz. During MRS acquisition, participants watched a nature documentary to minimise boredom and sleepiness. At each timepoint the participant watched the identical section of the documentary and was instructed to keep their eyes open throughout. This was monitored via an eye tracker, although no data were collected.

#### MRI analysis

##### fMRI analysis

Pre-processing for the fMRI analysis involved i) brain extraction using BET^3^ ii) motion correction with MCFLIRT^4^ iii) B0 distortion correction using the fieldmap, iv) spatial smoothing using a 5mm Gaussian kernel and v) high pass temporal filtering. FLIRT was used to register EPI images to the individual structural images (BBR method)^5^ and FNIRT was used to register to standard space.

##### MRS analysis

MRS data were analysed using a local version of Gannet 3.1. This tool is coded in MATLAB (The Mathworks, Inc) and uses the Optimisation and Statistics toolboxes and SPM8. Preprocessing steps involved: 3Hz exponential line broadening, phase and frequency correction, outlier rejection and zero-padding. The GABA+ peak at 3-ppm was then modelled as a five-parameter Gaussian model. GABA+ (GABA+macromolecules) and Glx (Glutamate+Glutamine) were calculated as a ratio to total creatine (tCr). The total creatine (tCr) integral of a two- Lorentzian model of creatine and choline metabolites in the OFF spectrum was used to calculate creatine.

#### Data quality

##### Exclusions

Behaviour: Participants were removed from behavioural analyses if they did not complete the task accurately. For this reason, R010 was removed due to issues with task understanding during data collection at the post-training visit.

MRS: As previously described (Willis et al, 2023), MRS data were excluded from analysis if 1) software failed to fit the data or 2) neurochemical concentrations exceeded 3SD from the mean. hMT+ data were excluded from R013 (hMT+), R010 (SMC) and R022 (hMT+; SMC).

fMRI: In order to be considered for further analysis, there needed to be sighted field activity in the occipital cortex to 50% or 100% in both pre-training and post-training session and in at least 3 of the 4 conditions. These criteria were set because this level of visual stimulation should consistently activate the occipital regions in the healthy hemisphere. Therefore, consistent lack of activity suggests a lack of attention or possibly a short sleep. Using these criteria, 3 participants were excluded: R010, R017 and R023, leaving n=15 for the pre- and post-training fMRI data. In addition, in the follow-up group, a further participant (R006) was excluded due to excessive head motion, leaving n=10.

Supplementary Table 2. Participants excluded from analyses and reasons

| **Participant** | **Excluded from:** | **Exclusion Reason:** |
| --- | --- | --- |
| R001 | All analyses | Did not complete training |
| R008 | All analyses | Did not complete training |
| R009 | All analyses | Did not complete training |
| R010 | Behaviour; fMRI; MRS SMC | Misunderstanding during task; Lack of sighted fMRI activity; MRS values exceeding 3SD from the mean. |
| R012 | All analyses | Did not complete training |
| R013 | MRS hMT+ | post-training visit exceeding 3SD from mean |
| R017 | fMRI | Lack of sighted fMRI activity |
| R022 | MRS hMT+ and SMC | post-training visit exceeding 3SD from mean |
| R023 | fMRI | Lack of sighted fMRI activity |

##### Voxel positioning

MRS data were collected from two key regions of interest. The primary region was situated using the T1-weighted structural image within the extrastriate visual motion area, hMT+. The voxel was aligned parallel to the lateral cortical surface and further refined based on activity observed during the functional motion localiser^6^. For the control sensorimotor voxel, it was placed on the ‘hand knob’ of the ipsilesional central sulcus, aligned parallel to the dorsolateral cortical surface. Data acquisition from each voxel took approximately 20 minutes, including the setup time. Due to the limitations of patient tolerance and time constraints, data from the contralesional hemisphere or additional brain regions could not be acquired. The average voxel placement for each timepoint and voxel is indicated in Supplementary Figure 1.

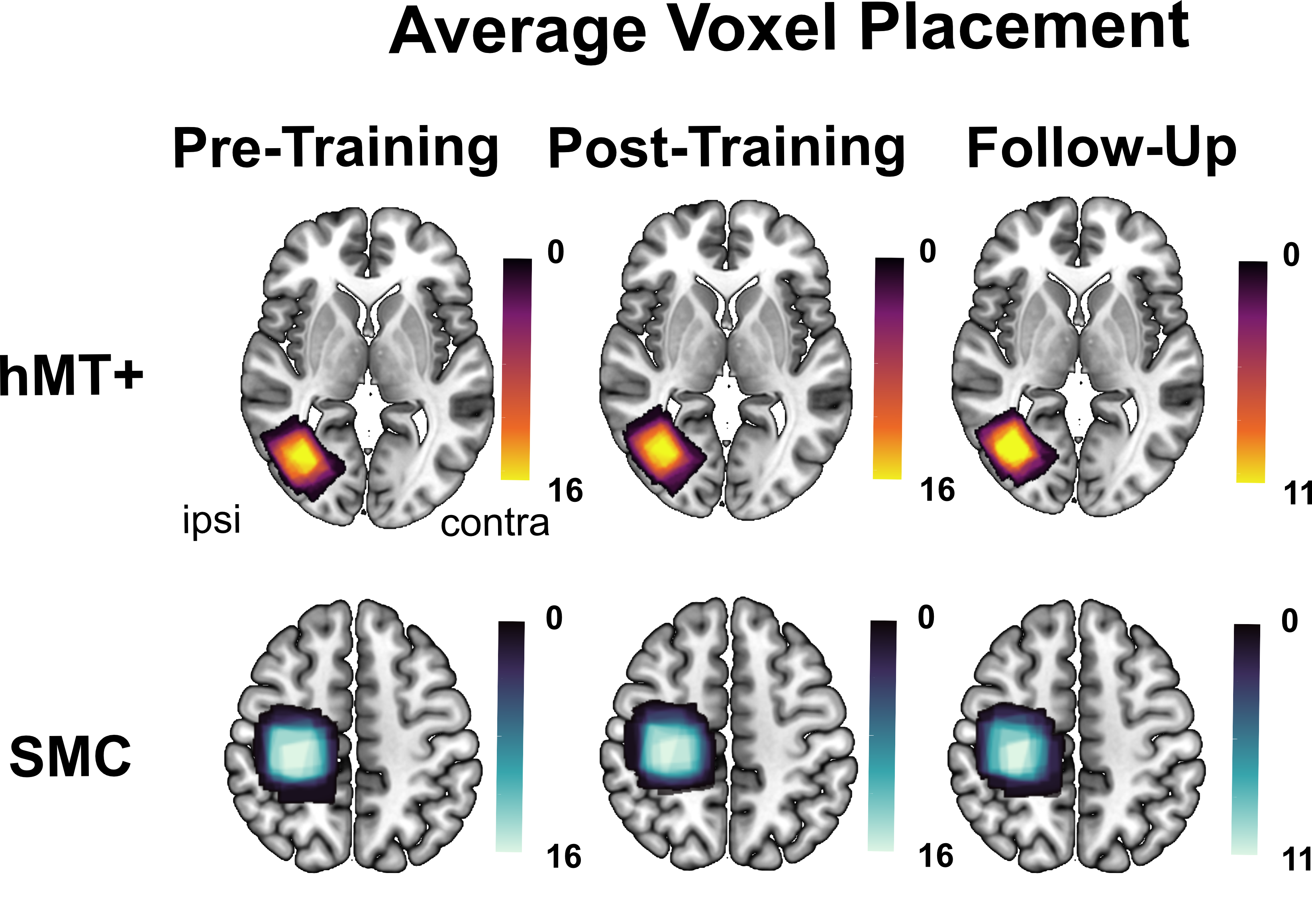

Supplementary Figure 1. Average voxel placement for each timepoint and voxel

To ensure the voxel was placed in similar locations at each timepoint, screenshots were taken during the pre-training scanning session. At the later time points (post-training, and follow-up), the research team (including trained radiographers) replaced the voxel using these screenshots. The overlap between the pre-training and post-training timepoints was calculated by registering the post-training and follow up voxels to the baseline using the FLIRT tool in FSL and then calculating the percentage overlap with the original voxel using the *fsl_maths* command. This analysis showed that between the pre- and post-training visits, the overlap of the voxels was on average 88.5% (IQR=10.1; range=74.16-99.49%) for hMT+ and 93.8% (IQR 14.9; range= 74.14-99.95%) for SMC. Between the pre-training and follow up visits, the overlap was on average 89.5% (IQR=7.15; range= 60.45-99.19) for hMT+ and 93.3% (IQR=9.89; range= 76.8 100%) for SMC. Based on this analysis, we are confident that the voxel was placed in roughly the same locations at each timepoint.

##### MRS Data quality

The Gannet output was visually inspected by two trained researchers for evidence of lipid contamination or poor water suppression. Fit error and full-width at half-maximum (FWHM) were calculated to determine whether they were similar across timepoints. Fit error and FWHM were compared between timepoints and between controls and stroke survivors (at the baseline visit) to determine whether data quality differed and could have explained the results. There was no significant difference in data quality across any of the comparisons (Supplementary Tables 3 and 4).

Supplementary Table 3. FWHM for GABA/tCr and Glx/tCr for each timepoint.

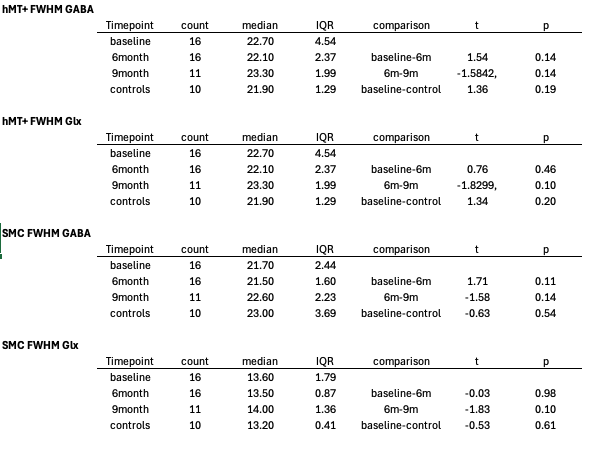

Supplementary table 4. Fit error for GABA/tCr and Glx/tCr for each timepoint.

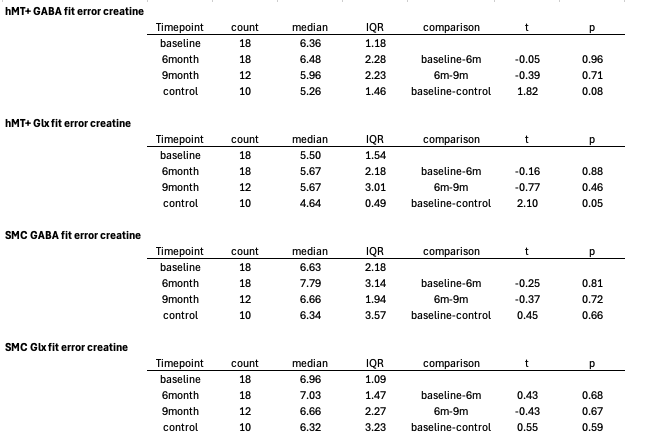

#### Additional results

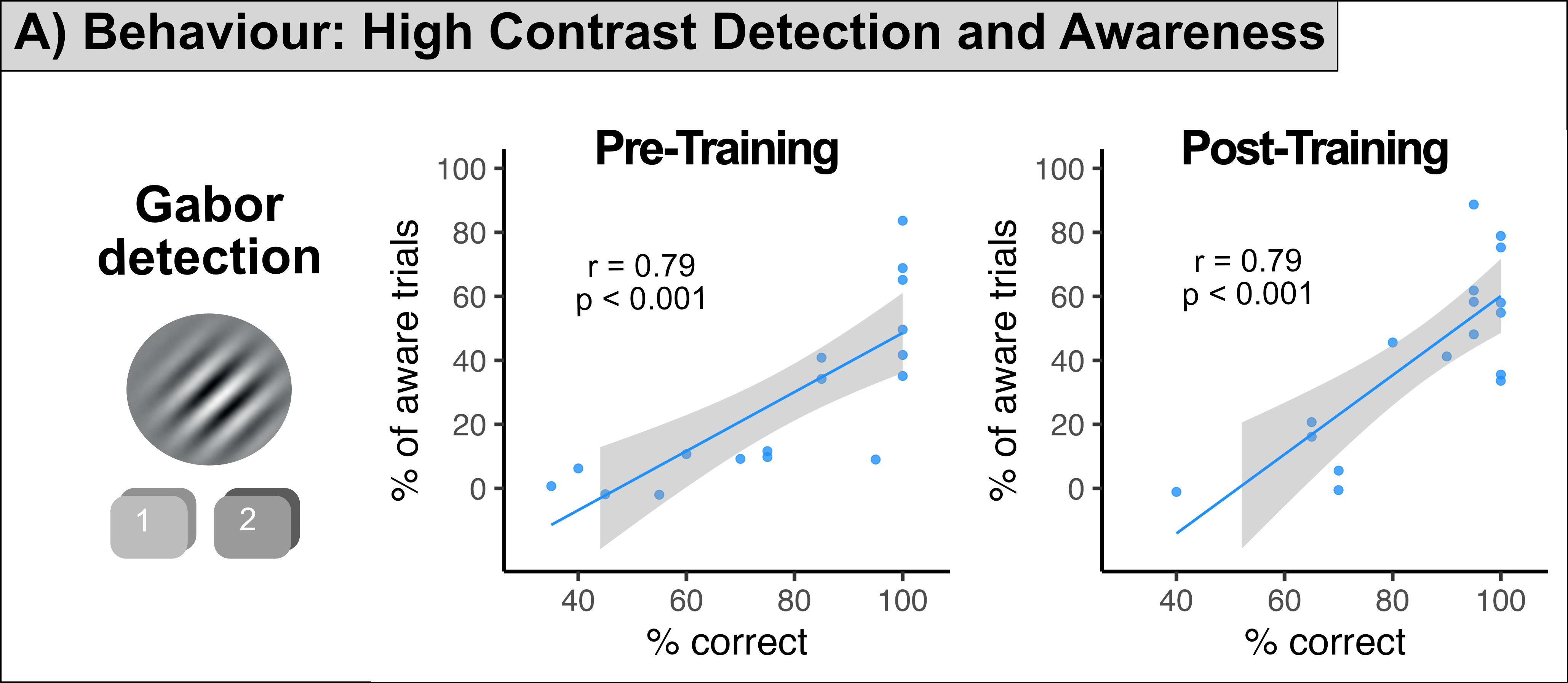

*Supplementary Figure 2.* Relationship between performance and awareness on the moving Gabor detection task at the pre- (left) and post-training (right) visits. There was a strong correlation between high contrast percentage correct and high contrast awareness at each timepoint.

##### Behavioural

To determine whether participant subjective awareness was related to accuracy in the task, we calculated the number of trials that were correct and aware, incorrect and aware, correct and unaware, and incorrect and unaware across the two timepoints. Average number of trials across the 50% and 100% contrast were calculated. We fitted a linear mixed model using the *lme4* package to predict percentage of correct trials with awareness and timepoint (formula: percentage of trials ~ awareness * timepoint + (1| Subj). The model included participant as a random effect. There was a significant main effect of awareness, *F*(1, 76) = 8.01, *p* = 0.006, indicating that accuracy differed between aware and unaware trials, with participants showing better performance when they reported being aware. There was no main effect of time, *F*(1, 76) = 0.31, *p* = 0.58, indicating that performance did not significantly improve with time. However, there was a significant interaction between awareness and time, *F*(1, 76) = 4.30, *p* = .042, suggesting that participants were more aware of the stimulus when giving correct answers after training.

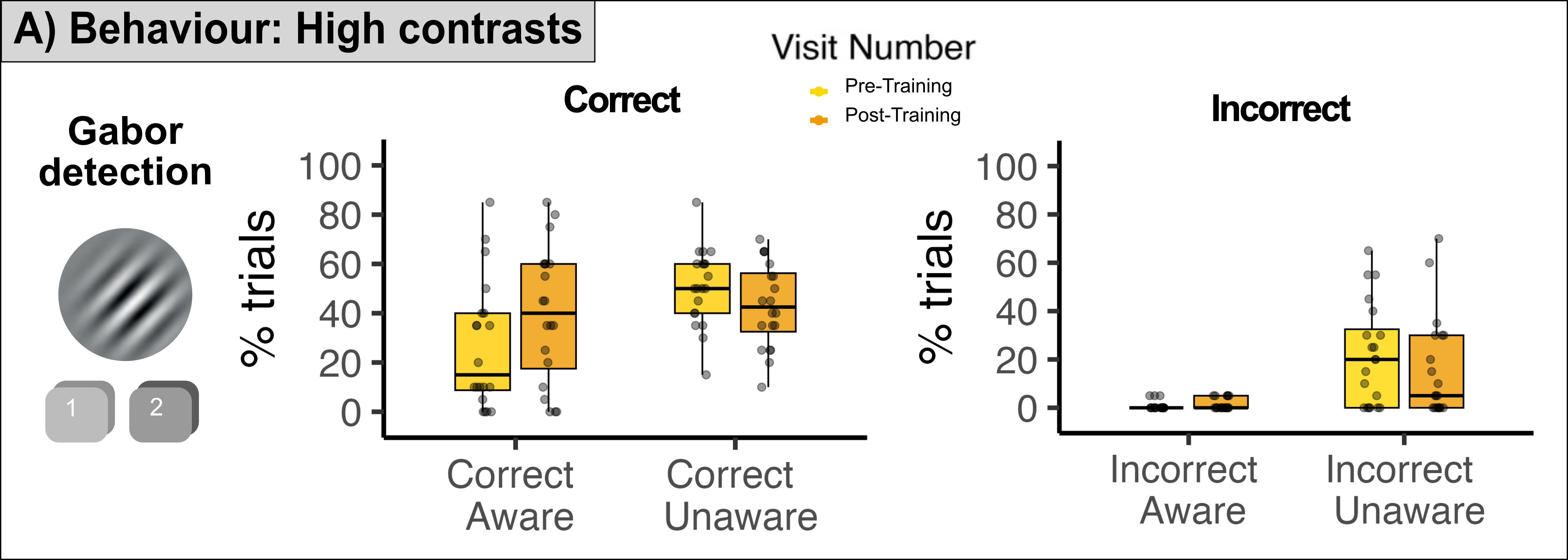

*Supplementary Figure 3.* Behavioural performance for high contrasts for aware and unaware trials. *Percentage of correct (left) and incorrect (right) trials is shown as a function of awareness (aware vs. unaware) and visit number (pre-training, post-training). Points represent individual participants, and boxplots indicate the median and interquartile range, with whiskers extending to the range of the data.*

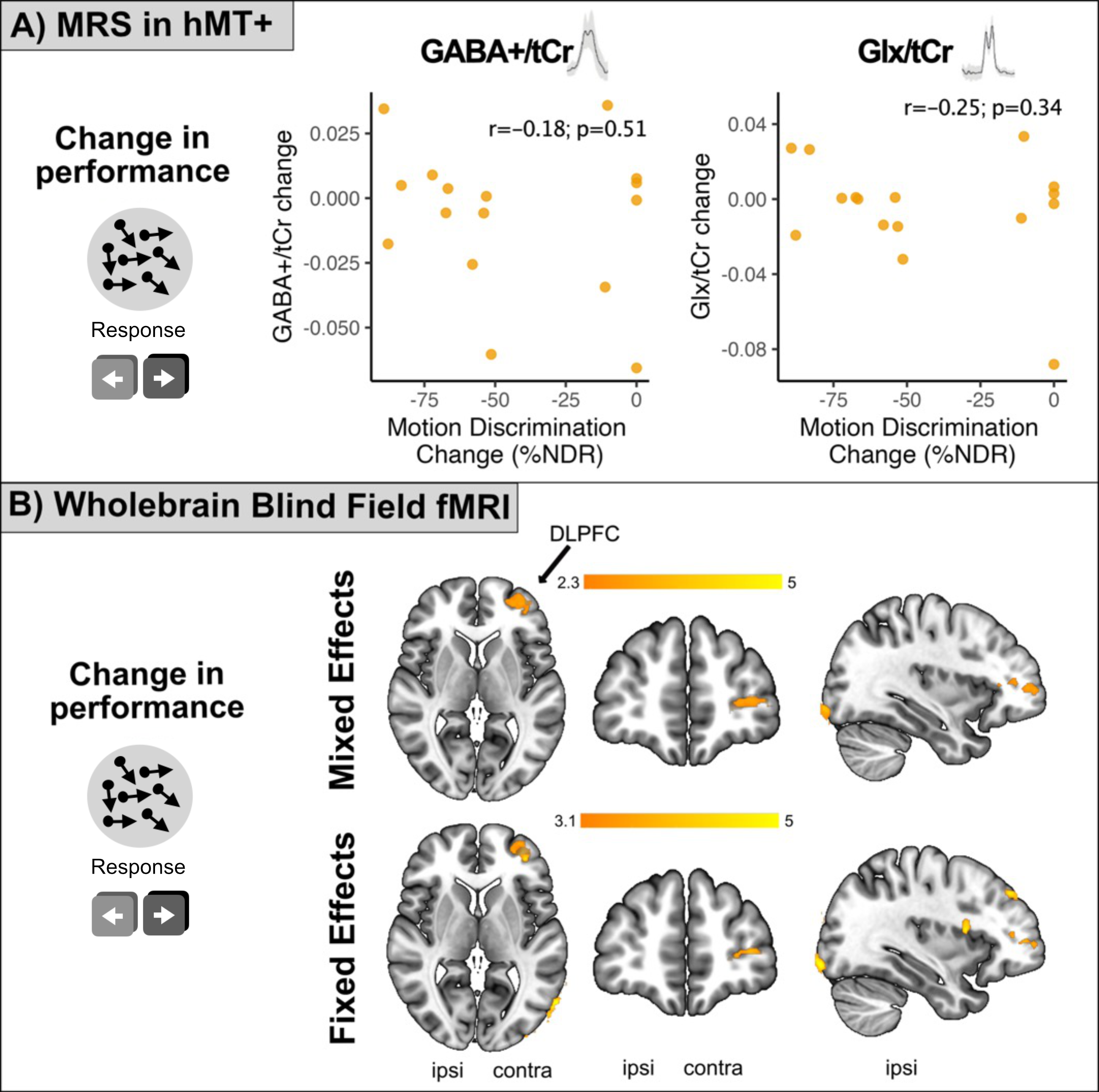

*Supplementary Figure 4.* Neural correlates of change in motion discrimination performance. A) Shows no relationship between the change in motion discrimination performance (normalised direction range thresholds) and GABA+/tCr or Glx/tCr concentration in hMT+. B) Shows the regions of the brain where the increase in performance was correlated with increased BOLD signal change to blind field stimulation. Both the mixed and fixed-effects analysis shows greater activity in the DLPFC in the hemisphere contralateral to the damage for those who show improvement in visual performance. “Ipsi” and “contra” indicate the ipsilesional and contralesional hemispheres respectively.

##### fMRI

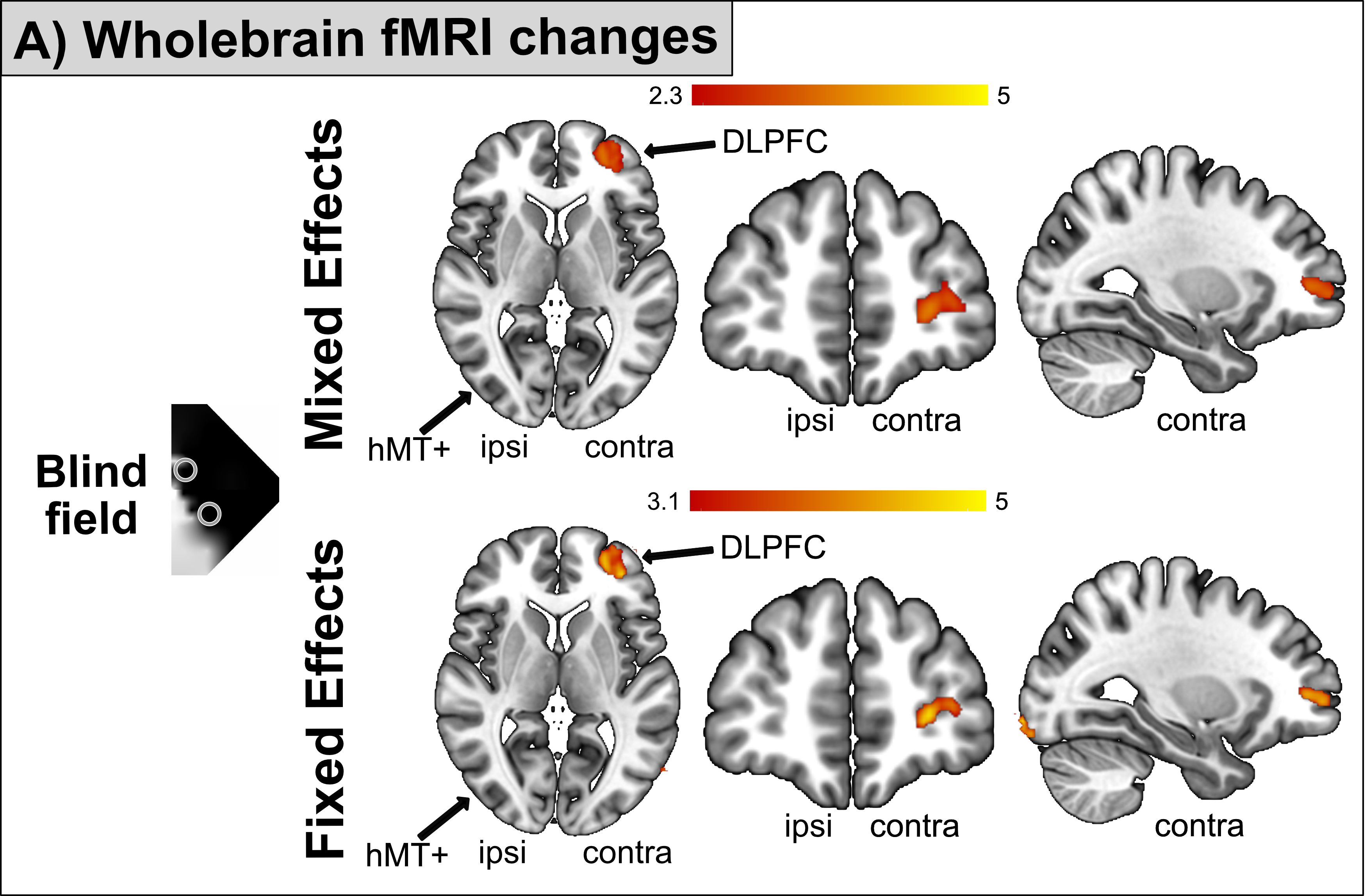

Supplementary Figure 5. The increase in neural activation to high-contrast stimuli in the post-training compared to the pre-training scan. Using both a mixed-effects and fixed effects analysis, the dorso-lateral prefrontal cortex showed a significant change to blind field stimulation, while no change in hMT+ was found. Ipsilesional (ipsi) and contralesional (contra) hemispheres are indicated.

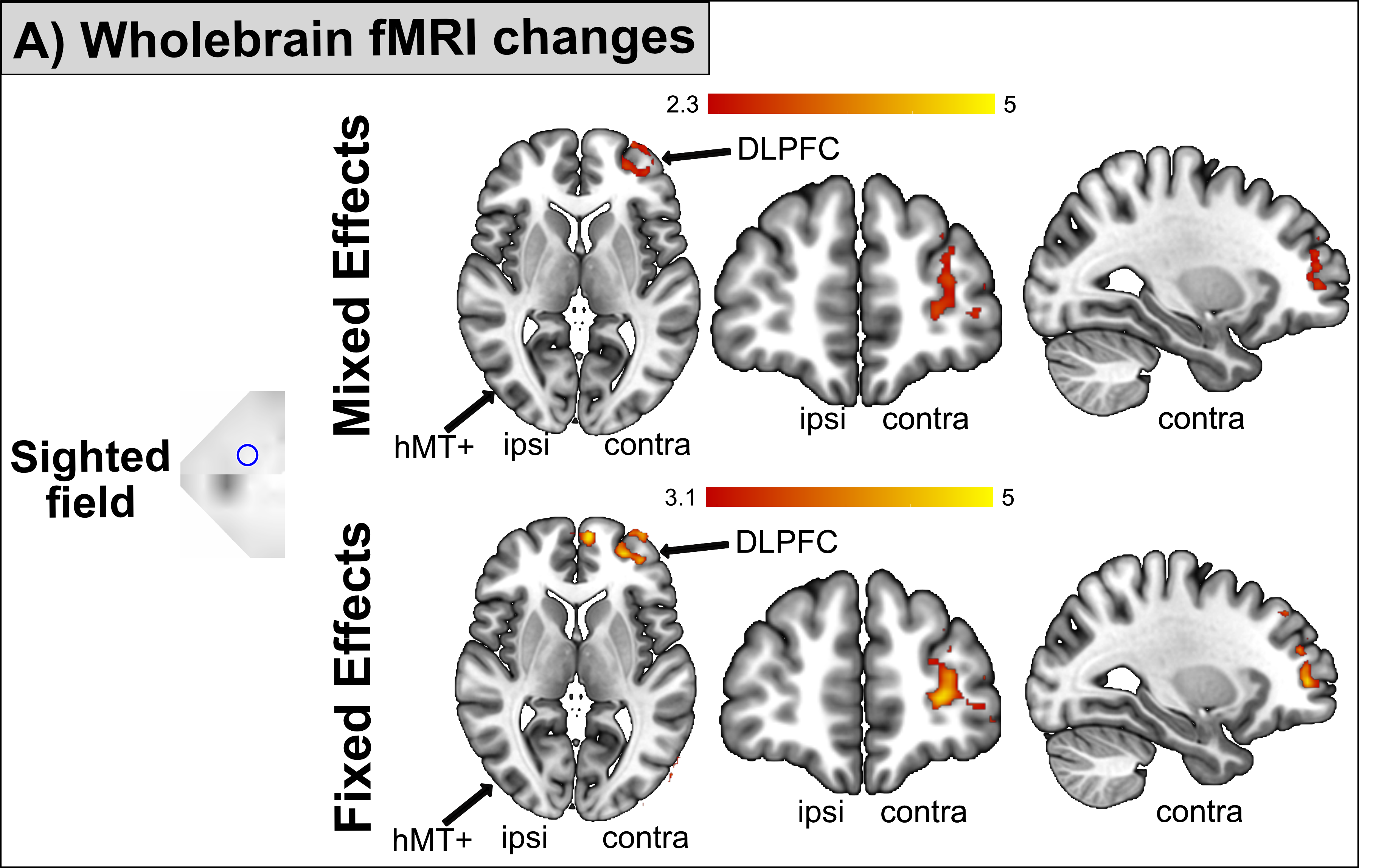

Supplementary Figure 6. The increase in neural activation to high-contrast stimuli presented to the sighted field between the pre- and post-training scan. Mixed-effects and fixed effects analysis both show activity in the dorso-lateral prefrontal cortex to sighted field stimulation, while no change in hMT+ was found.

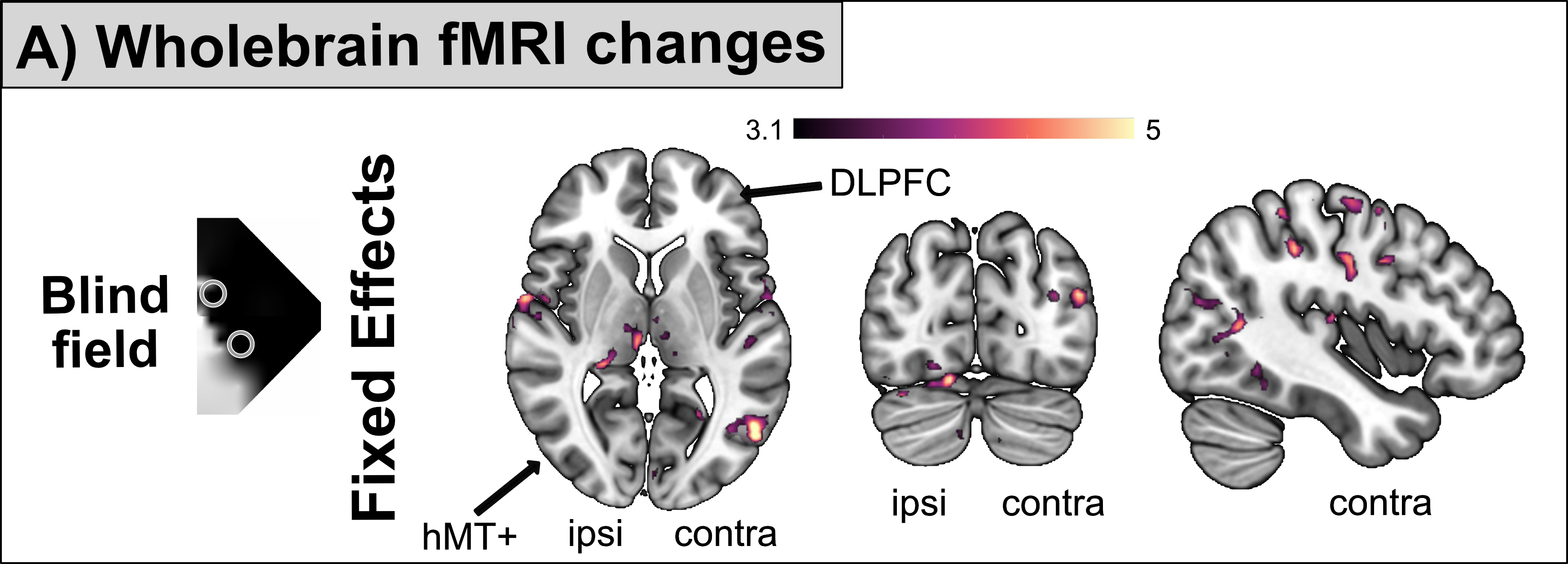

Supplementary Figure 7. Mixed effects analysis shows no changes in the neural activity between the post-training and follow-up fMRI scans. Fixed effects analysis (only considers changes in the occipital lobe) shows no reduction in neural activity between the post- and follow up sessions, but there was an increase in occipital activation. This activity shows increased activation in ipsilesional ventral visual cortex and an increased in contralesional hMT+ activation to blind field stimulation.

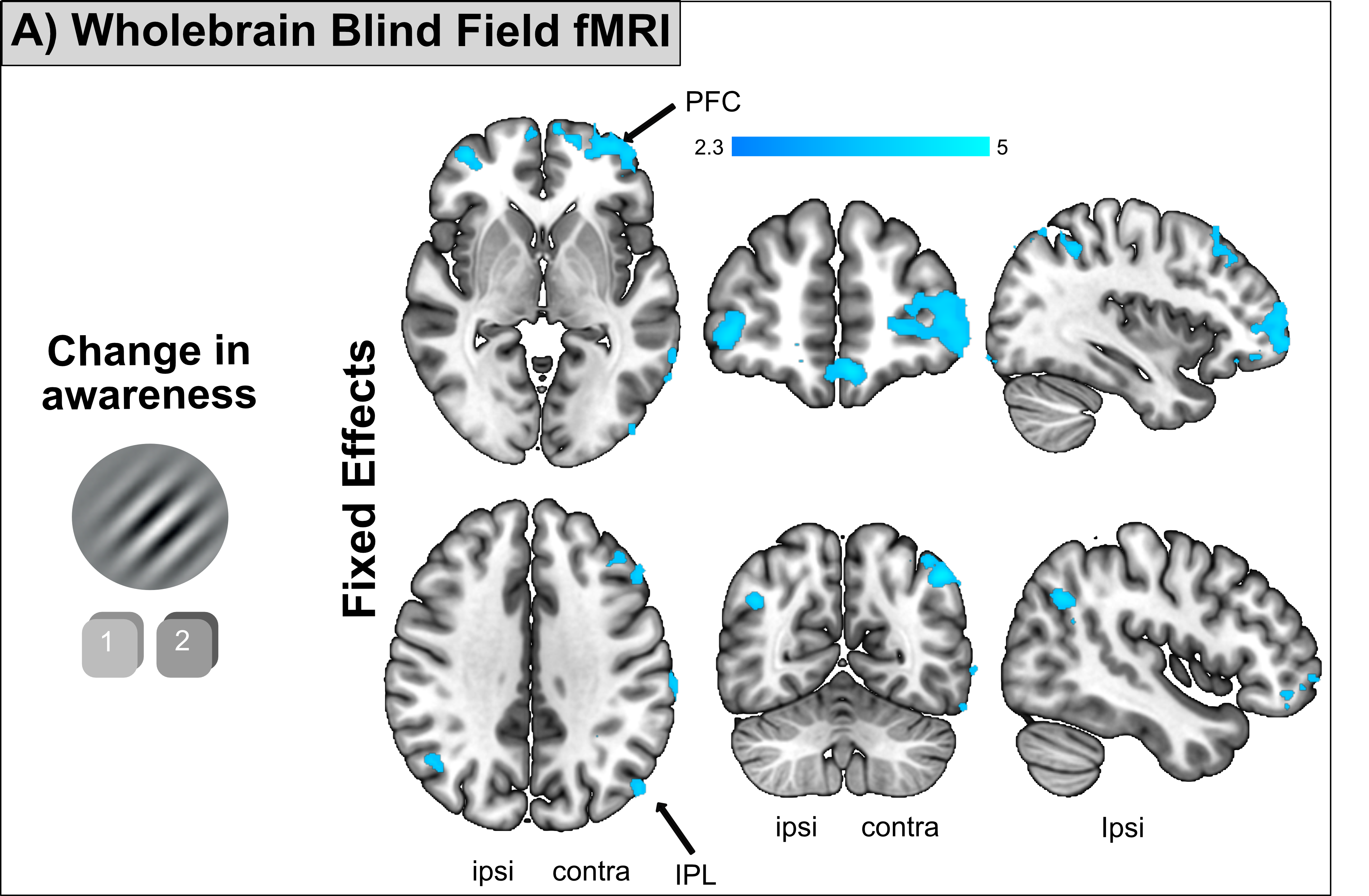

Supplementary Figure 8**.** Shows the regions of the brain where the increase in awareness was correlated with increased BOLD signal change across the group to blind stimulation after training using a fixed-effects analysis. Analysis shows greater activity in several areas compared to the mixed-effects analysis, including bilateral prefrontal cortex (area 46) and the inferior parietal lobule.
